# Biphasic acclimation for simultaneous wide-field fluorescent Ca^2+^ imaging and fMRI of awake mice

**DOI:** 10.1101/2025.10.27.684863

**Authors:** Francesca Mandino, Taekyung Kang, Xilin Shen, Corey Horien, Xenophon Papademetris, Stephen M Strittmatter, Evelyn MR Lake

## Abstract

Functional magnetic resonance imaging (fMRI) can be applied in mice and humans making it a key technology in translational neuroimaging research. Yet, most fMRI studies in rodents use anesthesia to limit subject motion and stress. This puts a hard boundary on the range of brain and behavioral states that can be studied in animals, and deviates from the near-universal practice of imaging people whilst awake. Recent years have seen a push towards the development of acclimation protocols for performing fMRI in mice without anesthesia, but the results have been mixed. In parallel, imaging mice without anesthesia has become routine for complementary neuroimaging methods including wide-field fluorescence calcium (WF-Ca^2+^) imaging. Our group works with a unique co-implementation of WF-Ca^2+^ and fMRI which enables the collection of complementary measures of brain function, but – until now – has necessitated the use of anesthesia. We present the first longitudinal protocol for simultaneous WF-Ca^2+^ and fMRI in awake mice. Our approach is comprised of a two-phase (biphasic) acclimation protocol that results in high-quality multimodal data. Comparisons of cortex-wide neuronal activity (WF-Ca^2+^ imaging) and blood-oxygen-level dependent fMRI signals between isoflurane-anesthetized and awake mice reveal both convergent and divergent patterns in brain function between modalities, and across time.

## 1. Introduction

Functional magnetic resonance imaging (fMRI) is a noninvasive tool for mapping large-scale brain networks that can be applied in both humans and animal models. These attributes make this modality a keystone technology within basic and translational neuroscience research (Mandino *et al*., 2019; Mandino *et al*., 2022; Pagani *et al*., 2023; Sheline *et al*., 2013; van den Heuvel *et al*., 2010; Van Essen *et al*., 2012; Xu *et al*., 2022). Yet, the blood-oxygen-level-dependent (BOLD) fMRI signal is incompletely understood and offers low spatiotemporal resolution, relative to other imaging modalities (Logothetis *et al*., 2004). In animal models, invasive methods can be applied in combination with fMRI to improve our collective understanding of brain function, and the BOLD signal (Lake *et al*., 2022). Although rare, due to the challenges of combining imaging methods with fMRI, much stands to be gained through the development of strategic simultaneous multimodal imaging approaches (Lake *et al*., 2020; Lake *et al*., 2022).

Wide-field fluorescence calcium (WF-Ca^2+^) imaging provides a cell-type specific measure of concert neural activity at high spatiotemporal resolution, relative to fMRI (Barson *et al*., 2020; Cardin *et al*., 2020; Lake *et al*., 2020; Mandino *et al*., 2025a; O’Connor *et al*., 2022; Vafaii *et al*., 2024). Although, the WF-Ca^2+^ imaging field-of-view (FOV) is limited to optically accessible tissue, this method provides sufficient overlap with fMRI for two complementary measures of large-scale activity patterns (Lake *et al*., 2022).

The first simultaneous acquisition of WF-Ca^2+^ and fMRI was described recently by our group (Lake *et al*., 2020; Mandino *et al*., 2025a; Vafaii *et al*., 2024). This work offered a unique framework for studying neurogliovascular signals and large-scale brain organization in anesthetized mice. The use of anesthesia was necessary due to the significant challenges of performing fMRI in awake animals – let alone the additional difficulties of performing simultaneous WF-Ca^2+^ imaging within the MRI magnet. For a comprehensive discussion, see our recent systematic review, “*Where do we stand on awake fMRI in mice?*” (Mandino *et al*., 2024). Among our suggestions for how a concerted effort can help establish standardized and validated approaches for acclimating mice to undergoing fMRI whilst awake, we called for more studies which explicitly compare data collected from awake and anesthetized animals given the surprisingly limited number of direct comparisons in the literature (Dinh *et al*., 2021; Fadel *et al*., 2022; Gutierrez-Barragan *et al*., 2022; Tsurugizawa *et al*., 2021). As such, the outcomes of this work are the introduction of a first-of-its-kind acclimation protocol for simultaneous WF-Ca^2+^ and fMRI in awake mice, an analysis of data quality, comparisons between data acquired from anesthetized and awake mice, as well as an analysis of state-related changes in functional connectivity. We chose to focus on network-related measures affected by state as access to these measures requires the large overlapping FOV afforded by both modalities (a unique feature of this multimodal implementation). Further, these measures are of central importance to fMRI analyses done on data acquired from human subjects, and have been shown to capture state-related differences in brain function.

Unlike in fMRI research, over the last decade it has become common to collect WF-Ca^2+^ imaging data from awake and behaving animals, e.g., (Barson *et al*., 2020; Cardin *et al*., 2020; Cramer *et al*., 2023; Zhao *et al*., 2021)(for a review, see (Ren *et al*., 2021)). This shift away from using anesthesia, within both WF-Ca^2+^ and fMRI applications, has in-part been motivated by a growing understanding of the influence anesthesia has on neurogliovascular function, as well as large-scale brain activity patterns (Barttfeld *et al*., 2015; Demertzi *et al*., 2019; Grandjean *et al*., 2014; Greene *et al*., 2023; Lake, 2022; Mandino *et al*., 2024; Slupe *et al*., 2018). Imaging awake animals also allows for the full-range of brain states to be observed alongside behavior and task performance. It is also worth noting that the vast majority of research on humans is conducted on awake subjects and that inter-species translation may be improved by performing like experiments in awake animals (Barson *et al*., 2020; Cardin *et al*., 2020; Lake *et al*., 2022; Mandino *et al*., 2024; Reimann *et al*., 2020).

The present work introduces and evaluates an experimental framework for acclimating mice to undergoing longitudinal, simultaneous WF-Ca^2+^ and fMRI whilst awake. The work leverages the large overlapping FOV afforded by these modalities to relate cortical neuronal activity – here, measured using a pan-neuronal fluorescence marker – and brain-wide BOLD-fMRI measures of canonical network organization. Guided by protocols developed for fMRI in awake mice (Mandino *et al*., 2024), we devised a biphasic approach comprised of an initial training where naïve animals are acclimated to undergoing imaging whilst awake in incremental steps, followed by a truncated ‘refresher’ training prior to subsequent imaging sessions. Subject motion, data exclusion, temporal signal-to-noise ratio (tSNR), and multimodal connectome specificity (Grandjean *et al*., 2023) (a measure of ‘biologically plausible’ connectivity patterns) are used to quantify multimodal data quality. Comparisons are made between data collected from awake and anesthetized mice. Differences in functional connectivity are quantified at the connectome, network, and brain region level. Convergent and divergent patterns are examined between imaging modalities and across time. These complementary quality control measures and functional connectivity analyses are chosen due to the novelty of the techniques, and their implementation in awake mice, as well as the inherent strengths of the modalities being used – specifically, their ability to access large-scale brain organizational patterns.

Taken together, our findings indicate that the biphasic acclimation protocol successfully prepared mice for undergoing simultaneous WF-Ca^2+^ and fMRI whilst awake. The multimodal data acquired from awake animals are high-quality based on current methods of evaluation. A range of differences in multimodal connectivity patterns are uncovered which span connectome, network, and regional levels. These show both convergent, as well as some divergent patterns across imaging modalities and time. Overall, this unique study lays a solid foundation for future studies aimed at dissecting large-scale brain function in awake mice using fMRI and multimodal imaging.

## 2. Methods

All procedures were approved by the Yale Institutional Animal Care and Use Committee (IACUC) and followed the National Institute of Health Guide for the Care and Use of Laboratory Animals. Both biological sexes were included. Mice were group housed in individually ventilated cages with access to food and water *ad libitum*. Animals were on a 12h light/dark cycle. Data from awake mice are compared with data from anesthetized animals. All data was acquired concurrently. MRI data from anesthetized mice has been previously published (Mandino *et al*., 2025b).

### 2.1 Surgical procedures for multimodal imaging

See **Fig. 1A** for an experiment timeline. Mice undergoing multimodal imaging (N=11) underwent two surgical procedures as described previously (Hamodi *et al*., 2020; Mandino *et al*., 2025b).

**Fig. 1.**
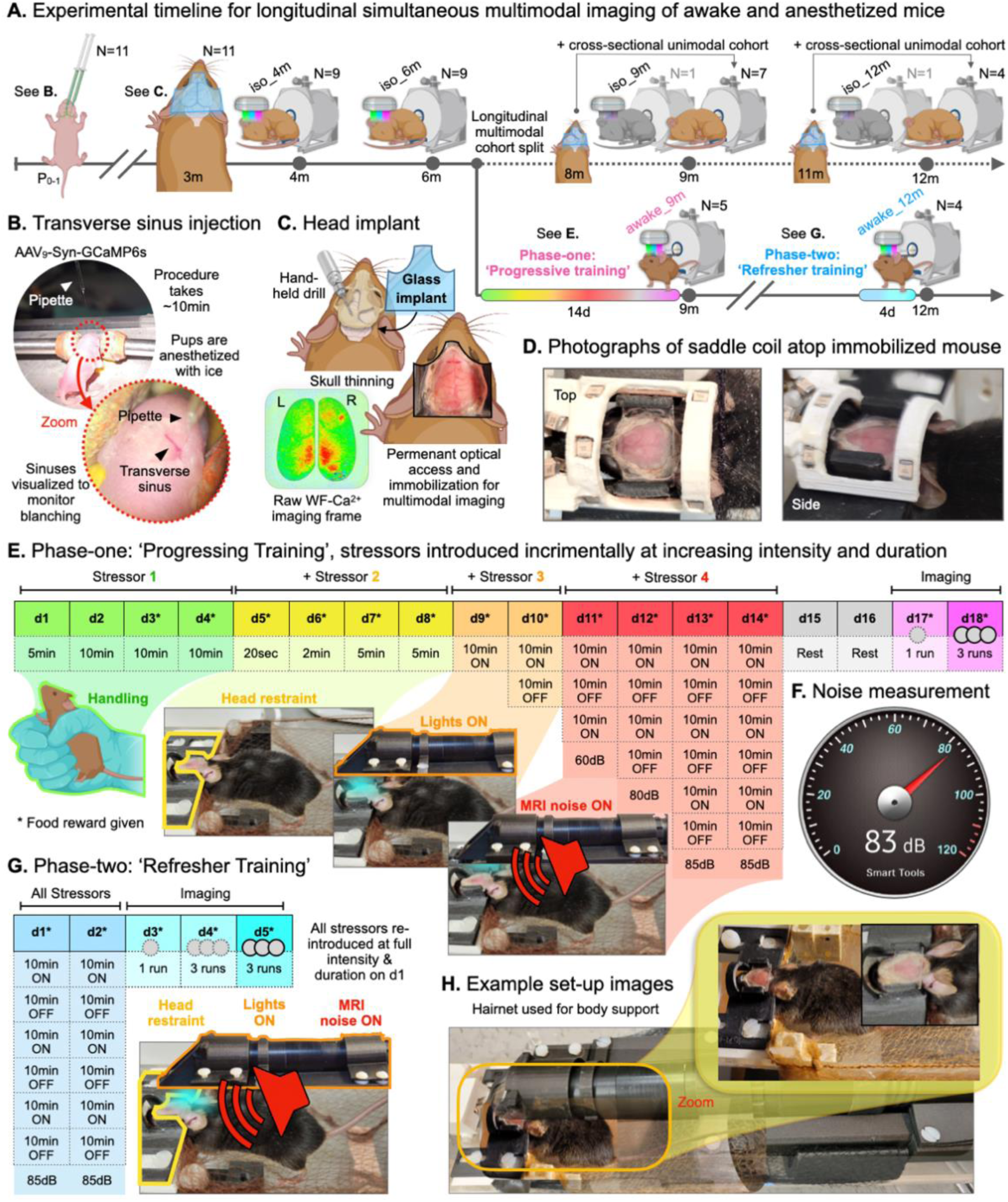
Experiment overview. **A.** Mice in the longitudinal multimodal imaging group (N=11) underwent transverse sinus injections (**2.1.1**), head-implant placement (**2.1.2**), and two multimodal imaging sessions at 4 and 6m under anesthesia (isoflurane) (**2.2**). Mice were then randomly assigned to an awake and anesthetized group. Animals in the awake group underwent two phases of acclimation training for awake multimodal imaging (**2.3**) and were imaged after each phase at 9 and 12m. Two additional cross-sectional unimodal (MRI-only) groups were added at the 9 and 12m imaging timepoints, after undergoing skull-thinning and head-plate application (**2.5**). Greyed-out mice indicate data excluded from all analyses. **B.** At birth, the viral vector AAV_9_.Syn.GCaMP6s.WPRE.SV40 was injected to induce pan-neuronal whole-brain fluorescence as described previously (Hamodi *et al*., 2020). **C.** At 3m, mice in the longitudinal multimodal imaging group underwent skull-thinning and head-implant placement. Mice in the unimodal (MRI-only) imaging groups also underwent head-implant placement one-month prior to imaging (**A.**). **D.** Photographs of the in-house built saddle coil for multimodal imaging. **E.** Protocol for acclimating mice to awake multimodal imaging (**2.3.1**). **F.** MRI-noise measurement example. Measured volumes were comparable between the real scanner and the mock set-up used for acclimation training (60-85dB). **G.** Protocol for re-acclimating mice to awake multimodal imaging (**2.3.2**). **H.** Additional photographs of a mouse being prepared for awake multimodal imaging.

#### 2.1.1 Transverse sinus injections of a viral vector for uniform whole-brain fluorophore expression

To elicit whole-brain Ca^2+^ indicator expression across neurons (pan-neuronal), transverse sinus injections of AAV_9_.Syn.GCaMP6s.WPRE.SV40 (Addgene plasmid #100843; http://n2t.net/addgene:100843; RRID:Addgene_100843) were administered at birth (P_0-1_) (Chen *et al*., 2013), **Fig. 1B**.

Pups were anesthetized using ice for 2–3min then transferred to a cool metal plate. A light microscope was used to visualize the transverse sinuses throughout the procedure. Sterilized fine scissors (Fine Science Tools, CA, USA) were used to make two small incisions (∼2mm) in the skin above each transverse sinus. A glass capillary tube (3.5’ #3-000-203-G/X, Drummond Scientific Co, PA, USA), pulled using a P-97 pipette puller (Sutter Instruments, CA, USA) was used for the injection. Pipettes were filled with mineral oil (M3516, Sigma-Aldrich, NY, USA) and attached to a Nanoject III (Drummond Scientific Co) and MP-285 micromanipulator (Sutter Instruments). Most of the mineral oil was ejected and replaced with vector solution. The pipette tip was lowered through the skull and into the lumen of the transverse sinus (300–400mm below the skull surface). With no delay, 2μL of vector solution was injected at a rate of 20nL/sec. The pipette was retracted after a 5sec delay, and the procedure repeated targeting the opposite hemisphere. Delivery was verified by the observed blanching of the sinus. With the two injections complete, the skin was sealed with VetBond, and the pup placed on a warming pad. The procedure typically lasted 10min. The total volume (titer of 1×10^13^vg/mL) of vector solution injected per pup was 4μL.

To minimize rejection by the mother, the whole litter was removed from the home cage together and kept on a warming pad while each pup underwent the procedure in sequence. Once all pups were alert, they were returned to the home cage and gently rubbed with home-cage bedding to mask foreign odors. This procedure resulted in cortex-wide GCaMP transfection and pan-neuronal expression by adulthood in 10/11 pups. No WF-Ca^2+^ imaging was performed in the 11^th^ animal due to hydrocephalus being observed in the MRI data (**2.6**).

#### 2.1.2. Skull-thinning and head-implant placement for optical access to the cortex and immobilization

At 3 months-of-age (m), animals were initially anesthetized with 5% isoflurane (70/30 medical air/O_2_) and secured in a stereotaxic frame (KOPF) using ear bars and an incisor bar. Isoflurane was then reduced to 2%. Eye ointment was applied; meloxicam (1-5mg/kg body weight) administered subcutaneously, and bupivacaine (0.1%, 0.04ml/5mg) injected locally at the incision site. Fur was removed and the scalp washed 3 times with betadine followed by ethanol (70%). The skin and soft tissue overlying the desired FOV was surgically removed and the skull cleaned and dried. Antibiotic powder (Neo-Predef) was applied to the incision site, and isoflurane reduced further to 1.5%. Skull-thinning of the frontal and parietal skull plates was performed with a hand-held drill (FST, tip diameters 1.4 and 0.7mm). Notably, the skull was thinned but remained intact, **Fig. 1C**. Superglue (Locite) was applied to the exposed skull, followed by transparent dental cement C&B Metabond (Parkell). A pre-cut (Neurotechnology Core, USA) all-glass head-plate was attached to the dental cement prior to solidification. Animals were allotted 3 weeks to recover before any imaging data were collected or acclimation procedures begun.

### 2.2 Data acquisition

The dataset is comprised of longitudinal simultaneously acquired WF-Ca^2+^ and fMRI data (N=11) collected using an apparatus and imaging protocol we have described previously (Lake *et al*., 2020; Mandino *et al*., 2025a; O’Connor *et al*., 2022; Vafaii *et al*., 2024) as well as cross-sectional unimodal (MRI-only) data (N=11) to offset animal attrition (**2.5**). Mice in the unimodal dataset underwent head-plate implantation but not viral vector transfection (**2.1**). Imaging parameters (for both modalities) do not vary across timepoints or between groups. Mouse body weight was monitored, during acclimation training for awake imaging, as a proxy for chronic stress (Mandino *et al*., 2024), **Fig. S1**. For mice that were imaged using anesthesia, a free-breathing low-dose isoflurane regimen (∼0.5-0.75%, in 50/50 medical air/O_2_) was used as we have described previously (Lake *et al*., ^2^02^0^; O’Connor *et al*., 2022; Vafaii *et al*., 2024). Mice that were imaged whilst awake underwent a biphasic acclimation training protocol (**2.3**). This included both a ‘Progressive Training’ phase (**2.3.1**), conducted prior to the first awake imaging timepoint, **Fig. 1E**, and a ‘Refresher Training’ phase (**2.3.2**), conducted prior to the second awake imaging timepoint, **Fig. 1G**.

#### 2.2.1 (f)MRI

Data were acquired on an 11.7T preclinical magnet (Bruker, Billerica, MA), using ParaVision v6.0.1. Body temperature was monitored (Neoptix fiber), maintained with a circulating water bath (36.6-37°C), and recorded (Spike2, Cambridge Electronic Design Limited). For acquisitions where anesthesia was used, breathing rate was measured using a foam respiration pad (Starr Life Sciences Corp., USA). During data acquisition, imaging and physiological measures were synchronized and recorded (Master-8 A.M.P.I., Spike2 Cambridge Electronic Design Limited).

*Structural MRI.* For image registration (Lake *et al*., 2020; Mandino *et al*., 2025a):

1. A multi-spin-multi-echo (MSME) image sharing the same FOV as the fMRI data. Acquired with a repetition time (TR) of 2500ms and echo time (TE) of 20ms, 28 slices, two averages, and resolution of 0.1×0.1×0.31mm^3^ (10min and 40sec),
2. A whole-brain isotropic 3D image using a MSME sequence, with 0.2×0.2×0.2mm^3^ resolution, TR/TE of 5500/20ms, 78 slices, and 2 averages (11min and 44sec), and
3. A fast-low-angle-shot (FLASH) time-of-flight (TOF) angiogram (FOV, 2.0×1.0×2.5cm^3^, covering the cortex), TR/TE of 130/4ms, and resolution of 0.05×0.05×0.05mm^3^ (18min).

### Functional MRI

Data were acquired using a gradient-echo echo-planar imaging (GE-EPI) sequence: TR/TE of 1.8sec/11ms, 334 volumes, 25 slices, and resolution of 0.31×0.31×0.31mm^3^ resulting in near whole-brain coverage (∼10min per run). For anesthetized imaging timepoints, three runs were acquired per timepoint (30min per mouse).

Following the ‘Progressive Training’ for awake imaging (**2.3.1**), two consecutive days of imaging were performed. On day-one, mice underwent a truncated protocol that included only one functional imaging run (10min per mouse). On day-two, three functional imaging runs were acquired akin to the anesthetized imaging protocol (30min per mouse), **Fig. 1E**. Following ‘Refresher Training’ (**2.3.2**), three consecutive days of imaging were performed which included one, three, and three functional imaging runs (10, 30, and 30min per mouse), **Fig. 1G**.

#### 2.2.2. WF-Ca^2+^ imaging

Data were acquired simultaneously with fMRI. The custom-built optical components which enable in-scanner WF-Ca^2+^ imaging were described by us previously(Lake *et al*., 2020; O’Connor *et al*., 2022; Vafaii *et al*., 2024). Our in-scanner WF-Ca^2+^ imaging set-up has a FOV of 1.4cm^2^ and collects optical images with a 25×25μm^2^ resolution. Data were recorded using CamWare version 3.17. As we have described previously(Lake *et al*., 2020; O’Connor *et al*., 2022; Vafaii *et al*., 2024), data were acquired using two wavelengths: violet (395/25, GCaMP-insensitive) and cyan (470/24, GCaMP-sensitive) interleaved (LLE 7Ch Controller, Lumencor) at 20Hz, to enable frame-by-frame background correction (inclusive of hemodynamic correction), and a resultant effective sampling rate of 10Hz. Both wavelengths have an exposure time of 40ms and are separated by 10ms of no illumination to avoid rolling shutter artifacts.

### 2.3 Biphasic training for awake multimodal imaging

#### 2.3.1 Phase-one: ‘Progressive Training’

Performed prior to the first awake multimodal imaging session. Four sources of stress (stressors) were introduced incrementally at increasing intensity and duration during a 14-day period, **Fig. 1E**. Maximum duration (60min) and intensity were reached on day-13 (and repeated on day-14).

Stressors:

1. Experimenter handling (performed exclusively on day-1 through -4),
2. Head immobilization (with supported body suspension, i.e., hammock),
3. Illumination (necessary for WF-Ca^2+^ imaging), and
4. MRI-noise (a recording of the scanner).

After 4-days of exclusive handling by a single (female) experimenter, mice were exposed to the remaining stressors over a 10-day period. This was done within a mock-scanner environment comprised of a semi-circular tray which held the animal, a head-plate fixation apparatus, the MR-coil, and the optical imaging equipment, as well as a black tube which slid overtop of the whole set-up to mimic the scanner bore. The tray, and associated parts, were the actual components used during subsequent imaging sessions. A food reward (condensed milk) was provided from day-3 onwards.

From the conclusion of handling (i.e., beginning of head immobilization on day-5), the mouse’s body was suspended in a ‘hammock-like’ bed constructed from a hairnet, **Fig. 1H**. This design grants body support without providing a surface for the mouse to effectively push against and likely helped preserve the integrity of the head-plate. One instance of head-plate detachment was observed (**2.5**).

For MR-noise, we used an application (Android ‘Sound Meter’, **Fig. 1F**) to measure true scanner noise (85–90dB). During training, the same application was used to monitor simulated scanning in the mock-scanner environment (60–85dB). Simulated scanning included a recording played back on speakers (Abramtek, China) placed next to the mock-scanner.

After 14-days of acclimation training, mice were given two days of rest (day-15 and -16), followed by two multimodal imaging sessions on consecutive days (day-17 and -18). On day-17, a truncated imaging protocol which included one functional run was performed (total scan time: 40min). On day-18, the full imaging protocol which included three functional runs was performed (total scan time: 60min). All anatomical imaging data needed for multimodal data registration (**2.4.3**) was collected at both imaging sessions.

#### 2.3.2 Phase-two: ‘Refresher Training’

For animals that have previously undergone Phase-one: ‘Progressive Training’ (**2.3.1**) and are not naïve to awake multimodal imaging. The protocol consisted of 2 days in the mock-scanner with all stressors reintroduced at full intensity and duration (60min), followed immediately by three multimodal imaging sessions on consecutive days, **Fig. 1G**. As done on the first imaging day following Phase-one, during the first imaging day following Phase-two (day-3), mice underwent a truncated imaging protocol which included one functional run (total scan time: 40min). The full imaging protocol, which included three functional runs (total scan time: 60min), was performed on the subsequent imaging days (day-4 and -5). All anatomical imaging data needed for multimodal data registration was collected at all imaging sessions (**2.4.3**).

Analyses were performed using data collected from awake mice on the last imaging day following Phase-one and Phase-two acclimation training. Specifically, data collected on day-18, following Phase-one, and day-5 following Phase-two.

### 2.4 Data preprocessing

#### 2.4.1 fMRI

Data preprocessing was performed as described previously (Mandino *et al*., 2025b), using RABIES v0.5.0 (Desrosiers-Grégoire *et al*., 2024). All EPI data from each mouse (at each imaging session) was averaged to create a mean image, corrected for intensity inhomogeneities, and non-linearly registered to the corresponding isotropic MSME anatomical image. This reduces susceptibility-induced distortions. Slice time correction was applied to the native space timeseries. To move the EPI data to common space, four transformations were applied: (1) motion correction, (2) mean fMRI → MSME, (3) MSME → within-dataset template, and (4) within-dataset template → out-of-sample template. We use an in-house out-of-sample template generated from N=162 datasets as described previously (Mandino *et al*., 2025b) which has been publicly shared (**2.4.3**). Transformations were concatenated then applied to each imaging frame. Data were resampled from the acquisition (0.31×0.31×0.31mm³ isotropic) to the template (0.2×0.2×0.2mm³ isotropic) resolution. Results were visually inspected using the RABIES-QC report (Rodent automated BOLD improvement of EPI sequences, quality control). Motion parameters, as well as the global signal, cerebral spinal fluid, and white matter (WM) signals were regressed, and framewise displacement (FD) computed. Frames with FD>0.075mm were scrubbed and interpolation used to bridge gaps (Desrosiers-Grégoire *et al*., 2024). Data were smoothed using a 0.4mm Gaussian kernel.

#### 2.4.2 WF-Ca^2+^ imaging

Data were preprocessed as we have described previously (Mandino *et al*., 2025a). Briefly, GCaMP-sensitive and -insensitive channels were smoothed with a 0.1mm sigma, and down-sampled by a factor of two in each spatial dimension. The GCaMP-insensitive channel was regressed from the GCaMP-sensitive channel (to denoise and remove hemodynamic contamination) and ΔF/F_0_ (F, fluorescence) computed for each run.

#### 2.4.3 Multimodal data registration

Data from both imaging modalities were filtered (0.008-0.2Hz) and thirty seconds discarded from the beginning and end of each run. This mitigates edge artifacts caused by temporal filtering in the fMRI data (Desrosiers-Grégoire *et al*., 2024) and eliminates the “photobleaching” artifact in the WF-Ca^2+^ imaging data (O’Connor *et al*., 2022). Given the greater temporal resolution of WF-Ca^2+^ imaging, an additional ‘fast’ frequency band was also analyzed (0.4-4Hz), as we have done previously (Mandino *et al*., 2025a; Vafaii *et al*., 2024).Timepoints censored based on MRI-measured FD were also removed from the WF-Ca^2+^ imaging timeseries. Thus, the data from both modalities were temporally matched. To spatially register the multimodal data, the WF-Ca^2+^ images were moved to the out-of-sample template using three transformations: two linear (1) WF-Ca^2+^ → Angiogram, and (2) Angiogram → MSME and one nonlinear (3) MSME → in-house template, as described by us previously (Lake *et al*., 2020; Mandino *et al*., 2025a; Vafaii *et al*., 2024). This procedure utilizes specialized tools developed for this purpose within BioImage Suite, BIS (https://bioimagesuiteweb.github.io/webapp/), see **Fig. 2** in (Mandino *et al*., 2025a). Via the BIS platform, the tools, atlases (**2.6.3**), and the in-house out-of-sample template have been made freely available.

**Fig. 2.**
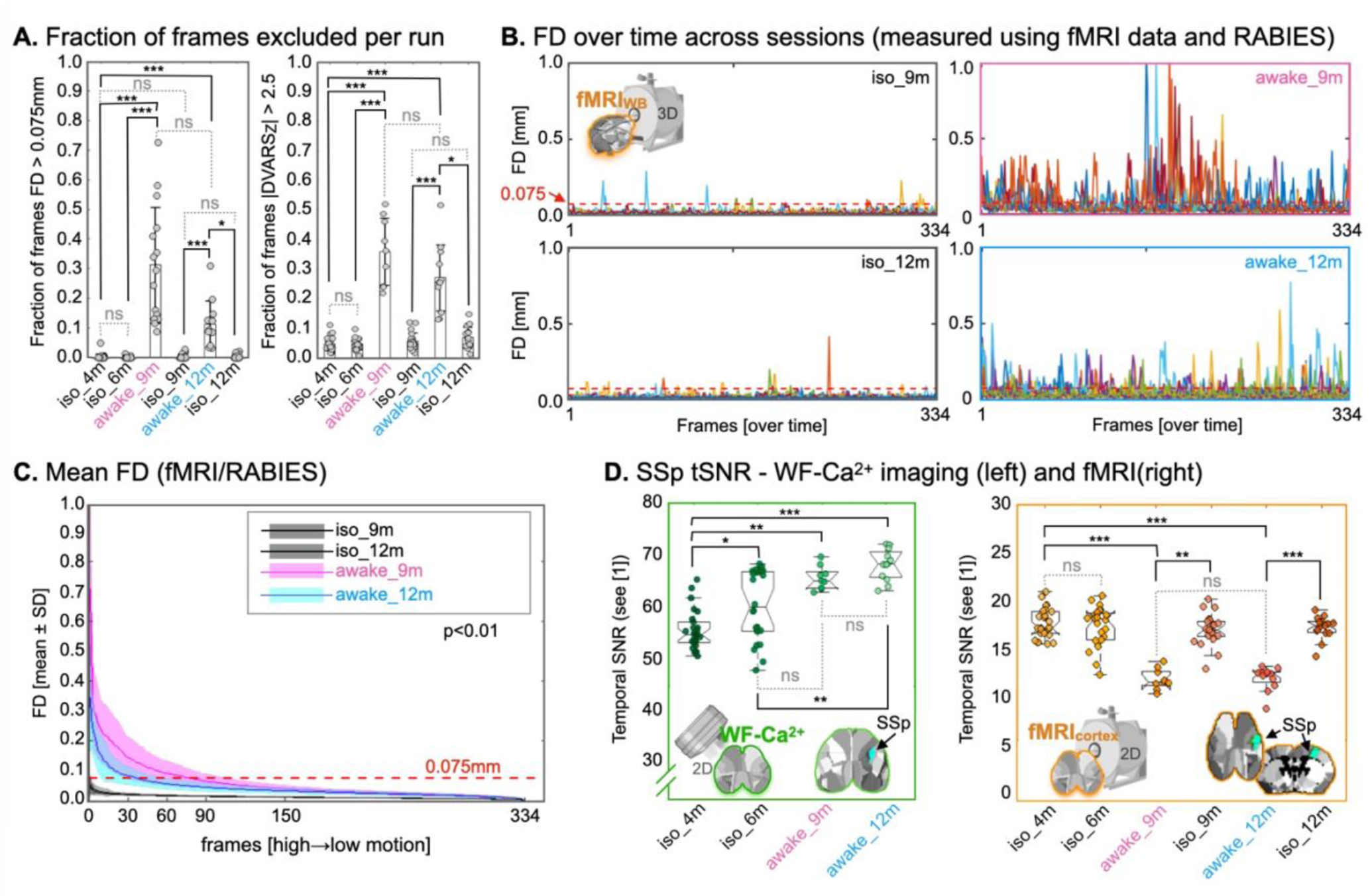
Multimodal signal quality measures. **A.** Based on fMRI data, using RABIES, the fraction of imaging frames per run excluded based on motion FD>0.075mm (left), or |DVARS_Z_|>2.5 (right). More imaging frames were excluded, based on either criterion, from awake relative to anesthetized imaging sessions. **B.** Also based on fMRI data, FD estimated by RABIES, over time. Different runs/mice are color-coded. The 0.075mm FD threshold for frame exclusion is indicated by a red dashed line. More motion was evident in awake relative to anesthetized imaging sessions. A reduction in motion was seen following the second, relative to the first, awake imaging session. **C.** FD (mean ± SD), measured using fMRI data and RABIES, sorted from highest to lowest. A significant cross-session effect was recovered (area under the curve tested with ANOVA and pairwise Bonferroni correction). **D.** Temporal SNR (tSNR) in a representative cortical region of interest (ROI), right primary somatosensory, upper limb (SSp-ul), computed for each run – after exclusion criteria were applied (**2.5**) – using WF-Ca^2+^ imaging (green, left) or fMRI (orange, right) data. Diverging patterns between imaging modalities were observed. Corrected p-values are displayed: *p<0.05, **p< 0.01, ***p<0.001.

### 2.5 Exclusion criteria

N=11 pups underwent transverse sinus injections to elicit fluorophore expression (**2.1.1**). None were rejected by mothers or excluded based on an absence of sinus blanching (necessary for good transfection) (Hamodi *et al*., 2020). Of these, N=2 were excluded due to hydrocephalus at 4m. Between the 6 and 9m imaging timepoints, N=2 mice died due to complications with housing. Before the 9m imaging timepoint N=1 was euthanized due to an eye infection that did not result from the head-plate. Remaining mice were randomly allocated to an awake (N=5), and anesthetized (N=1) multimodal imaging group. Between the 9 and 12m imaging timepoints, N=1 was excluded from the awake group due to poor head-plate integrity.

To augment the number of mice in the anesthetized group at the 9 and 12m imaging timepoints, MRI data was collected from two additional cohorts: N=7 at 9m, and N=4 at 12m (total N=11), after skull-thinning and head-plate application. As these mice did not undergo viral vector transfection to elicit fluorophore expression, no WF-Ca^2+^ imaging data were acquired from these animals. All mice underwent head-plate implantation for head immobilization (**2.1.2**). Given the lack of WF-Ca^2+^ imaging data collected from anesthetized mice at the 9 and 12m imaging timepoints, these data were excluded from all analyses. From awake mice, only the n=3 runs acquired on the final day following each phase of training were analyzed (**2.3**).

Cohort:

1. 4m, N=9 (n=27 runs) anesthetized multimodal datasets,
2. 6m, N=9 (n=27) anesthetized multimodal datasets,
3. 9m, N=5 (n=15) awake multimodal, and N=7 (n=21) MRI-only anesthetized datasets,
4. 12m, N=4 (n=12) awake multimodal, and N=4 (n=12) MRI-only anesthetized datasets.

No mice were excluded during Phase-one: ‘Progressive Training’ (**2.3.1**) or Phase-two: ‘Refresher Training’ (**2.3.2**) due to weight loss, **Fig. S1**. At 6m, data from one mouse was excluded due to fMRI imaging artifacts. At 9m, data from one mouse in the MRI-only anesthetized group was excluded based on image registration failure. At the 4 and 6m imaging timepoints, no runs were excluded due to motion (>1/3 imaging frames <0.075mm FD) or the RABIES DVARS_Z_ (derivative of timecourses’ variance) criterion (|DVARS_Z_|>2.5, (Desrosiers-Grégoire *et al*., 2024)) which identifies imaging frames containing artifacts caused by motion as well as other sources. At the 9m imaging timepoint, 5 runs from awake mice were excluded due to FD or DVARS_Z_. These came from two mice, thus all the data from one mouse was excluded. In addition, one run was excluded from a third mouse due to WF-Ca^2+^ imaging artifacts. At the 12m imaging timepoint, no datasets or runs were excluded. Additional dataset information is summarized in **Table S1**.

Final dataset:

1. 4m, N=9 (n=27 runs) anesthetized multimodal datasets,
2. 6m, N=8 (n=24) anesthetized multimodal datasets,
3. 9m, N=4 (n=9) awake multimodal, and N=6 (n=18) MRI-only anesthetized datasets,
4. 12m, N=4 (n=12) awake multimodal, and N=4 (n=12) MRI-only anesthetized datasets.

For the multimodal datasets, if data was excluded due to criteria applied to one modality (e.g., motion measured in the fMRI timeseries, or imaging artifacts in one or the other modality) the corresponding data in the second modality was also excluded. Thus, all the multimodal data are paired.

### 2.6 Data quality metrics

#### 2.6.1 Temporal SNR

Functional MRI tSNR was computed using RABIES (Desrosiers-Grégoire *et al*., 2024) and [1] where *μ_t_* is the mean signal intensity (SI) of a voxel over the course of a run and *σ_t_* is the SD of the voxel’s SI over the same period. Similarly, the WF-Ca^2+^ imaging tSNR was computed using [1] in MATLAB (v2021b) where *μ_t_* is the mean SI of a pixel within the frame range 1,000-1,500 (50sec extracted from the middle of each run).

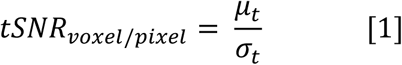

#### 2.6.2 Motion

Motion in the fMRI data was estimated using RABIES while BIS (and MATLAB as a comparison) were used to estimate motion in the WF-Ca^2+^ imaging data. When the 6 motion parameters, available from fMRI/RABIES, were considered independently, it was clear that the vast majority of subject motion was in the up/down or dorsal/ventral direction, **Fig. S2B**. Given that WF-Ca^2+^ imaging offers a 2D view from above, it follows that these data were “blind” to these movements, **Fig. S2C**. Still, data excluded based on motion in the fMRI timeseries were also excluded from the WF-Ca^2+^ imaging dataset (**2.5**) (more below).

### 2.7 Functional connectivity

#### 2.7.1 Brain atlases

Brain regions or nodes were based on the Allen Institute Common Coordinate Framework Reference Atlas (CCFv3) (Wang *et al*., 2020) as described by us previously (Mandino *et al*., 2025a). Results were generated using a 3D (whole-brain, fMRI), 166-node, or 2D (cortical, multimodal), 46-node, bilaterally symmetric version of the atlas. Brain regions included in the 2D version were based on coverage in the WF-Ca^2+^ imaging FOV, **Fig. S3**; brain regions included in the 3D whole-brain version were based on EPI slice package coverage, **Fig. S3**.

#### 2.7.2 Connectomes and network definitions

To compute connectomes, WF-Ca^2+^ or fMRI signals within each region were averaged, and the inter-regional Pearson’s correlation computed, and Fisher transformed. Connectomes were computed for each run using data residing in the in-house template space, as described by us previously (Mandino *et al*., 2025a; Mandino *et al*., 2025b). Regions were subdivided into seven whole-brain networks (Gutierrez-Barragan *et al*., 2022; Mandino *et al*., 2025a; Xu *et al*., 2022), two of which were well represented within the cortex, **Fig. S3**. To provide a more fine-grained analysis of cortical network organization, these were further subdivided into four sub-networks: motor (MO), somatosensory/auditory (SS/AUD), visual (VIS) and default-mode network (DMN).

#### 2.7.3 Multimodal “specific connectivity”

Motivated by recent work which uses relative connectivity strengths to gauge ‘biologically plausible’ connectivity patterns (Grandjean *et al*., 2023), connectivity between the right and left SSp, SSp_R_ ↔ SSp_L_, were plotted against connectivity between SSp_R_ and the right anteromedial visual area (VIS), SSp_R_ ↔ VIS_R_. In this framework, data with high connectivity strength (correlation>0.1, z-scores) in SSp_R_ ↔ SSp_L_ and coincident low connectivity strength (correlation<0.1, z-scores) in SSp_R_ ↔ VIS_R_ are considered desirable. These data were plotted alongside a reference distribution created from randomly chosen homotopic node pairs, and inter-hemispheric node pairs with n=1,000 iteration for each.

### 2.8 Analyses and statistics

Significance is indicated as follows: *p<0.05, **p<0.01, ***p<0.001. All statistical tests were computed using MATLAB version R2021b.

#### 2.8.1 Multimodal signal QC

Unless stated otherwise, for all analyses assessing motion and tSNR, Kruskal-Wallis tests (MATLAB, v.R2021b) were applied to assess differences across imaging sessions. For any significant effects (p<0.05), post hoc pairwise multiple comparisons were performed and Bonferroni correction applied. Equality of variance was tested with Levene’s test (MATLAB, v.R2021b). Body weight was measured with repeated measure ANOVA (*fitrm* and *ranova*, Greenhouse-Geisser correction, MATLAB).

#### 2.8.2 Connectome specificity

Distributions were generated by randomly selecting pairs of homotopic or non-homotopic nodes from each hemisphere. Comparisons were made using a two-sample Kolmogorov-Smirnov (KS) test, with Bonferroni correction applied to account for multiple comparisons. The proportion of runs falling within the boundaries defined for “specific” connectivity (**2.7.3**) were computed as a fraction of the total number of runs for each condition at each imaging timepoint.

#### 2.8.3 Connectivity differences

##### Edge-wise

Connectivity strength between pairs of brain regions within the cortex (main figures) or across the whole-brain (**Supplementary Material**) were compared across conditions (anesthetized vs. awake states) using two-sample t-tests (*ttest2*, MATLAB, v.R2021b) for each edge, followed by Benjamini-Hochberg correction to control for multiple comparisons. T-values are displayed without a threshold (bottom left half) and with a threshold for statistical significance applied (top right half), **Figs. 4, S7-10**.

##### Cross-network

For both WF-Ca^2+^ and fMRI, differences in connectivity between data collected from anesthetized and awake mice was computed (ΔConnectivity) and a t-test (*ttest2*, MATLAB) applied to identify differences of ΔConnectivity between imaging modalities (Benjamini-Hochberg), **Fig. 4B**.

##### Within network

Changes in connectivity strength were assessed between data collected from anesthetized mice at the 4m timepoint and data collected from awake mice at the 9 or 12m imaging timepoints, independently, **Figs. 4C, S8** (analyses including both awake sessions are described below). Mean intra-hemispheric and inter-hemispheric functional connectivity was calculated for each functional network by averaging pairwise connectivity values within the corresponding network. Each imaging run constituted one observation, with multiple runs acquired from the same mouse retained in the analysis. To account for repeated measurements (intra- and inter-hemisphere, multiple networks, two states) acquired from the same mouse across multiple imaging runs, network-level connectivity values were analyzed using a linear mixed-effects model (*fitlme*, MATLAB R2021b), with state (awake vs. anesthetized), hemisphere (intra-vs. inter-hemispheric), and network included as fixed effects and mouse included as a random intercept. Significant interactions were further investigated by fitting separate linear mixed-effects models for each network and hemisphere, testing the effect of state while including mouse as a random effect. Non-parametric permutation tests were also used to evaluate shuffled-labels (n=1,000 iterations) versus true measures, **Fig. S11**.

##### Cross-modal correlation

Two-sampled t-tests were used to compare intra-versus inter-network correlation strengths and a one-sampled t-test against zero to identify significant coupling within each imaging session was applied, with Bonferroni correction (**Fig. 5**).

##### Longitudinal analyses

To assess *longitudinal* changes in functional connectivity across imaging sessions as well as state, edge-wise connectivity values were analyzed using linear mixed-effects models (*fitlme*, MATLAB R2021b), **Fig. S10**. Connectivity strength between pairs of brain regions was assessed at 4, 6, 9, and 12m in both awake and anesthetized conditions. Of note, no data for WF-Ca^2+^ anesthetized sessions was available at 9 and 12m (see **2.5**), so this analysis could only be completed for fMRI data. A linear mixed-effects model was fit with age (4, 6, 9, or 12 months), state (awake vs. anesthetized), and their interaction included as fixed effects, and mouse included as a random intercept to account for repeated measurements across imaging sessions. Statistical significance was evaluated separately for the main effects of age, state, and their interaction. Resulting p-values were corrected for multiple comparisons across all edges using the Benjamini-Hochberg false discovery rate procedure.

## 3. Results

At birth, mice underwent transverse sinus injections of AAV9.Syn.GCaMP6s to achieve pan-neuronal cortex-wide calcium indicator expression (Hamodi *et al*., 2020), followed by skull-thinning and head-plate implantation at 3m, as described previously (Mandino *et al*., 2025a) (**2.1**). This preparation enables head fixation and permanent optical access to the cortical surface for simultaneous WF-Ca^2+^ and fMRI (**Fig. 1A-C**). Mice underwent a longitudinal multimodal imaging protocol comprised of two imaging sessions under anesthesia (at 4 and 6m) and two whilst awake or anesthetized (at 9 and 12m), **Fig. 1A**.

Prior to being imaged whilst awake, mice underwent acclimation training. The biphasic acclimation protocol, consisted of both a progressive phase in advance of the first awake imaging session (at 9m) (**2.3.1**) and a refresher training phase prior to the second awake imaging session (at 12m) (**2.3.2**), **Fig. 1E-G**. Unimodal data (fMRI-only) were acquired from age-matched subjects, at 9 and 12m, under anesthesia to offset attrition in the second cohort (**2.5**). Body weight, signal QC metrics, and functional connectivity analyses follow. All p-values are corrected for multiple comparisons (**2.8**).

### 3.1 Longitudinal multimodal imaging of awake mice does not cause weight loss

Animal weight, measured daily during acclimation and imaging, was used as a proxy for significant prolonged stress (Mandino *et al*., 2024). Body weight was unaffected during both training phases and following awake imaging sessions, **Fig. S1**. While body weight does not capture subtle or acute stress responses, these results indicate that the acclimation and imaging procedures implemented here did not induce overt or sustained chronic stress. No statistically significant differences were found across training, for mice that were imaged whilst awake (progressive phase: p=0.08/0.28, males/females; refresher: p=0.64/0.5, males/females). Further, body weight of animals that underwent multimodal anesthetized sessions showed comparable results to awake animals (**Fig. S1**).

### 3.2 Multimodal signal quality

Fraction of data excluded, motion, and tSNR were quantified and compared across imaging timepoints. Functional MRI data were preprocessed using RABIES (**2.4.1**)(Desrosiers-Grégoire *et al*., 2024). The WF-Ca^2+^ imaging data were background corrected, smoothed, and down-sampled, before ΔF/F₀ was computed, as we have described previously (**2.4.2**)(Mandino *et al*., 2025a). Both WF-Ca^2+^ and fMRI data were censored based on fMRI-derived motion traces (FD<0.075mm)(Desrosiers-Grégoire *et al*., 2024). The multimodal data were spatially registered using anatomical landmarks, as described previously (**2.4.3**)(Lake *et al*., 2020).

#### 3.2.1 Awake mice moved more than anesthetized mice, but less after Phase-two: ‘Refresher Training’

The fraction of imaging frames excluded from anesthetized animals was low, stable across time, **Fig. 2A**, and in-line with previous work (Grandjean *et al*., 2020; Mandino *et al*., 2024), **Fig. S4A**. As expected, a greater fraction of imaging frames was excluded in the awake group, based on motion and |DVARS_Z_| >2.5 (Desrosiers-Grégoire *et al*., 2024), relative to anesthetized animals at both awake imaging timepoints (Mandino *et al*., 2024). Overall, there was a significant session effect on fraction of frames excluded based on FD and |DVARS_Z_| (Kruskal-Wallis, p<0.001, for both), whereby both awake sessions resulted in a significantly higher number of frames excluded, compared to anesthetized sessions (p<0.001, for both).

To quantify the amount of data retained after motion censoring, we summarized the number of remaining frames for functional connectivity analyses across imaging sessions (**Table S2**). As expected, anesthetized sessions retained nearly the full acquisition (mean retained frames: 312–320 of 334), whereas awake sessions retained fewer frames (214.9 ± 38.2 at 9 months; 244.3 ± 37.3 frames at 12 months), corresponding to a reduction in the number of censored frames following the second phase of acclimation (120.1 ± 38.2 vs. 90.7 ± 37.3). Despite the higher degree of censoring in awake imaging, the remaining data substantially exceeded the duration required for functional connectivity analyses, and the reduction in censored volumes after the refresher training is consistent with the improved motion profiles shown in **Fig. 2B&C**. Accordingly, between the first and second awake imaging sessions, the amount of motion was reduced (area under the curve 12 vs. 9m, p=0.0029), **Fig. 2B&C**. In general, motion in data acquired from both awake and anesthetized mice at all timepoints was in-line with previous work (Mandino *et al*., 2024), **Fig. S4**. Notably, motion observed in the fMRI data was predominantly along the dorsal– ventral axis (**Fig. S2B**) and was not reflected in the simultaneously acquired WF-Ca^2+^ imaging data (**Fig. S2C**), consistent with modality-specific sensitivity to motion-related signal changes.

#### 3.2.2 Multimodal tSNR showed divergent patterns across modalities and dependence on whether data were acquired from awake or anesthetized mice

The WF-Ca^2+^ imaging data showed a significant increase in tSNR across imaging sessions, **Fig. 2D**, with reduced variance at later (awake) imaging timepoints (χ², p<0.001, Levene’s test, p=1.802e-06). Specifically, WF-Ca^2+^ imaging tSNR in a representative example cortical region (primary somatosensory, upper limb, SSp-ul) was lower in 4 or 6m anesthetized mice compared to 9 or 12m awake mice (p<0.01, for both), respectively. On the other hand, fMRI tSNR in the same example cortical region, SSp-ul **Fig. 2D**, and other brain regions **Fig. S5** (Grandjean *et al*., 2020), was lower in awake relative to anesthetized mice (p<0.01), while fMRI tSNR variance decreased slightly in awake sessions compared to 4 and 6m anesthetized imaging sessions (p=0.04). The fMRI data (whole-brain) also showed the expected tSNR dependence on brain region (lower values near ear canals and in WM), **Fig. S5**, and was in-line with previous work (in anesthetized mice), in terms of expected range for this modality, based on multi-center data (Grandjean *et al*., 2020).

### 3.3 Multimodal “specific connectivity”

Connectivity between the right and left primary somatosensory, mouth, (SSp-m, abbreviated SSp for visualization purposes), SSp_R_ ↔ SSp_L_, were plotted against connectivity between SSp_R_ and the VIS_R_, SSp_R_ ↔ VIS_R_, to gauge ‘anatomically plausible’ connectivity patterns (**2.7.3**). This was motivated by previous work (Grandjean *et al*., 2023), which asserts — based on viral tracing experiments available from the Allen Institute (Atlas, 2004; Wang *et al*., 2020), **Fig. 3A** — that coincident high connectivity strength (correlation>0.1, z-scores) in SSp_R_ ↔ SSp_L_, i.e. regions anatomically connected, and low connectivity strength (correlation<0.1, z-scores) in SSp_R_ ↔ VIS_R_, i.e. SS vs. regions that do not show structural connections, should be recovered in ‘high quality’ data. To the best of our knowledge, this is the first application of this framework in WF-Ca^2+^ imaging data, **Fig. 3B**, and the first application in simultaneously acquired WF-Ca^2+^ and fMRI data.

**Fig. 3.**
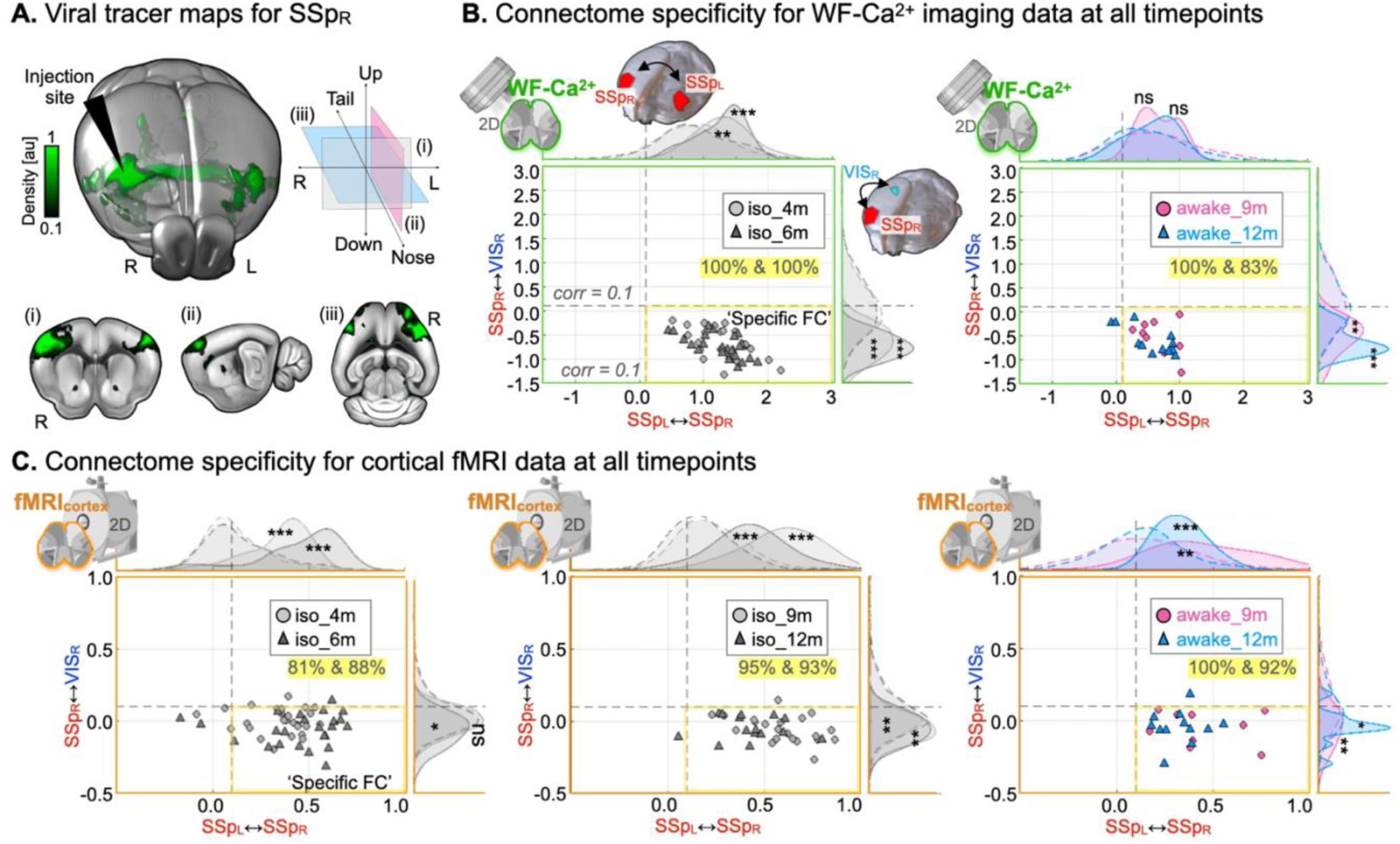
Multimodal “specific connectivity”. **A.** Viral tracer map showing the physical connections of primary somatosensory area, mouth, SSp_R_, based on viral vector injection (ID: 114290938, Allen Brain Connectivity Atlas) overlayed on an MRI template (average_template_50.nrrd, https://download.alleninstitute.org/informatics-archive/current-release/mouse_ccf/average_template/). The histology-based anatomical connection shown in this experiment motivates the “specific” functional connectivity framework for high versus low connectivity strengths SSp_R_ ↔ SSp_L_ and SSp_R_ ↔ VIS_R_, respectively. **B.** Scatterplots showing “specific” functional connectivity (**2.7.3**) using WF-Ca^2+^ imaging data. Data acquired from anesthetized mice are plotted in grey (left), while data acquired from awake mice are color coded (right). In the “specific” connectivity framework, high (correlation>0.1, z-scores) SSp_R_ ↔ SSp_L_ (x-axis) versus low (correlation<0.1, z-scores) SSp_R_ ↔ VIS_R_ (y-axis) connectivity strength should isolate ‘high-quality’ datasets in the lower right quadrant (highlighted in yellow). Above and to the right of the scatterplots, density distributions display kernel-smoothed observed (solid lines) and reference (dashed lines) distributions. Here, the reference distributions were generated from randomly selected intra- and inter-hemispheric node pairings (**2.7.3**). The fraction of runs identified as ‘high-quality’ are listed below the legend in each plot (highlighted in yellow). **C.** As in (**B.**) for fMRI data. Corrected (Bonferroni) p-values displayed: *p<0.05, **p< 0.01, ***p<0.001.

#### 3.3.1 Ample “specific connectivity” recovered in multimodal data acquired from anesthetized and awake mice

In the majority of runs (>81%), the fMRI data showed the anticipated ‘Specific’ pattern: high SSp_L_ ↔ SSp_R_ and low, or absent, SSp_R_ ↔ VIS_R_ connectivity, **Fig. 3C** (data points appearing in the quadrant outlined in yellow). This indicated that, within this framework, these data would be considered ‘high-quality’. Relative to the reference distributions, which we chose to define as other inter- or intra-hemispheric node pairs (**2.7.3**), anesthetized mice (grey) showed greater SSp_L_ ↔ SSp_R_ connectivity than other homotopic node pairs (p<0.001), and lower, or absent, SSp_R_ ↔ VIS_R_ connectivity relative to other intra-hemispheric node pairs (p<0.05, with the exception of the 4m timepoint, not surviving Bonferroni correction). Similarly, fMRI data from awake mice showed greater SSp_L_ ↔ SSp_R_ connectivity than other homotopic node pairs (p<0.01), and lower connectivity between SSp_R_ ↔ VIS_R_ compared to nulls (p<0.05).

The same framework was applied to the simultaneously acquired WF-Ca^2+^ imaging data, **Fig. 3B**. In both anesthetized imaging sessions (grey), SSp_R_ ↔ SSp_L_ connectivity was greater than other inter-hemispheric node pairs (p<0.01). Interestingly, every run (100%) of WF-Ca^2+^ imaging data, relative to fMRI runs, were considered ‘high-quality’. However, the WF-Ca^2+^ imaging data from awake mice — which had higher tSNR than the anesthetized data (**Fig. 2D**) — showed a slightly lower fraction of ‘high-quality’ runs at the 12m timepoint (83%). Furthermore, neither awake imaging timepoint showed significant differences from the null distribution for the SSp_R_ ↔ SSp_L_ axis. No runs (data) were excluded from further analyses based on whether the anticipated connectivity pattern considered in this subsection was observed.

### 3.4 Connectivity differences between awake and anesthetized mice

Next, we examined whether connectivity patterns show convergent or divergent organizational patterns that depend on brain state (awake vs. anesthetized) and imaging modality. Connectivity differences between data collected while mice were awake, at 9 and 12m, and data collected while mice were anesthetized, at 4 and 6m, were computed (t-test, awake subtract anesthetized), and displayed as t-value maps. Averaged connectomes are shown in **Fig. S3&S6**.

Cortical regions, visible with both WF-Ca^2+^ and fMRI, are highlighted as main findings. Results from higher-frequencies, using the WF-Ca^2+^ imaging data, are shown in **Fig. S7B**. Overall, no significant differences were observed between the fMRI-matched and higher frequency WF-Ca^2+^ imaging data – in-line with our previous work (Mandino *et al*., 2025a; Vafaii *et al*., 2024). Results from fMRI-only whole-brain analyses are shown in **Fig. S8-10**. Results which use the 4m imaging timepoint, as a “baseline”, are shown in **Fig. 4&S8**. When the 6m (anesthetized) imaging timepoint was used, in-place of 4m, very similar results were obtained, **Fig. S9**. Finally, also using only fMRI data (**2.5**), a linear mixed model which considered whether mice were awake or anesthetized and mouse age, was generated, **Fig. S10**.

**Fig. 4.**
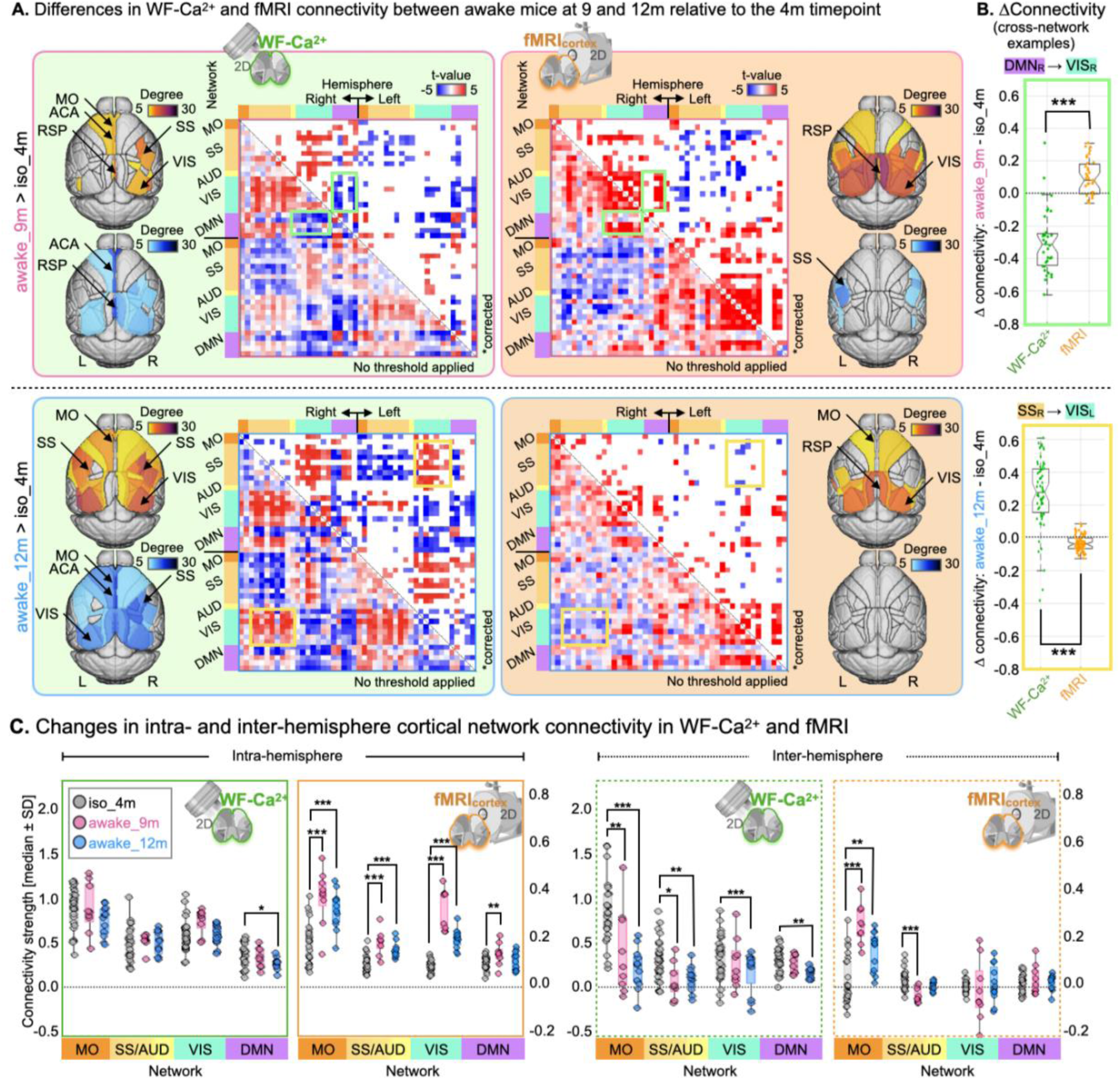
Differences in cortical WF-Ca^2+^ and fMRI connectivity between anesthetized and awake mice. **A.** Differences in WF-Ca^2+^ (left, green) and fMRI (right, orange) connectivity between the 9 (top, magenta outline) and 12m (bottom, cyan outline) awake imaging timepoints and the 4m (anesthetized) imaging timepoint. Results from a whole-brain analysis of the fMRI data are included in **Fig. S8**. An analysis which used the 6m (anesthetized) imaging timepoint, in-place of 4m, showed similar results, **Fig. S9**. Here, cortical regions, appearing in both imaging modalities **Fig. S3**, are shown. Matrices show significant connectivity differences (p<0.05) in the upper right half, and uncorrected differences in the bottom left half. Node-degree maps generated from corrected results appear to the right and left of each matrix (with a minimum of 5 edges as a threshold). Positive t-values (red, hot colors) indicate greater connectivity strength in awake mice. Negative t-values (blue, cool colors) indicate greater connectivity strength in anesthetized mice. **B.** Examples of cross-networks connectivity differences between awake and anesthetized mice that were different between imaging modalities (p<0.05, Benjamini-Hochberg, *ttest2*, MATLAB). Networks are indicated in (**A.**) by color coded boxes. **C.** For *a priori* functional networks (**2.7.2**, **Fig. S3**), connectivity strength (median ± SD) intra- (left, solid border) and inter-hemisphere (right, dashed border) at each imaging timepoint. Differences in connectivity assessed between 4m and either 9m (magenta) or 12m (cyan) awake sessions, respectively, with a linear mixed model each; plotted in the same graph for visualization purposes. Corrected p-values: *p<0.05, **p< 0.01, ***p<0.001. Spatial resolution: raw WF-Ca^2+^ = 25×25 μm^2^; raw fMRI = 310×310×310 μm^3^; Temporal resolution: raw WF-Ca^2+^ = 10Hz; raw fMRI = 0.55Hz.

#### 3.4.1 Widespread differences in cortical connectivity between awake and anesthetized mice were observed with both imaging modalities

At both the 9 and 12m awake imaging timepoints, relative to the 4m (anesthetized) imaging timepoint, WF-Ca^2+^ and fMRI showed widespread differences in cortical functional connectivity (**2.7**), **Fig. 4A**.

With fMRI at the 9m imaging timepoint, stronger connectivity was observed in awake relative to anesthetized mice in motor (MO), somatosensory (SS), retrosplenial (RSP), and visual (VIS) areas. Alongside these observations, weaker connectivity, in awake relative to anesthetized mice, was observed predominantly in inter-hemispheric somatosensory connections. At the 12m imaging timepoint similar trends were recovered in the fMRI data, but the effects were noticeably diminished and less widespread. As an additional control analysis, all runs across all sessions were truncated to the shortest retained scan length across imaging sessions before recomputing functional connectivity differences. This produced connectivity patterns that were highly consistent with those obtained using the full retained timeseries (**Fig. S7A**), indicating that the reported connectivity differences were robust to differences in retained timeseries length following motion censoring.

Conversely, the WF-Ca^2+^ imaging data showed an overall increase in connectivity differences that were more widespread at the 12 relative to the 9m imaging timepoint. These observations were all well replicated when the faster frequency band, accessed with WF-Ca^2+^ imaging, was utilized (**Fig. S7B**), or the 6m (anesthetized) imaging timepoint was used in-place of the 4m imaging timepoint, **Fig. S9**. Age-matched comparisons between awake and anesthetized groups (e.g. 9m awake vs. 9m anesthetized) — which were only possible with fMRI data (see **2.5**, and **Table S1**) — showed similar findings (**Fig. S9C**).

#### 3.4.2 Brain regions showed convergent and divergent changes in connectivity across modalities and imaging timepoints in awake relative to anesthetized mice

Brain regions (or nodes) showing changes in connectivity in awake relative to anesthetized mice in WF-Ca^2+^ and/or fMRI using data from the 9 and/or 12m imaging timepoints were identified using the corrected matrices in **Fig. 4A** (upper right halves). Common edges, across imaging modalities and timepoints, were identified by taking the product of binarized matrix pairs (after statistical thresholds for significance were applied, **2.8**). Node-degree, and bilateral symmetry, were used to identify regions showing common or divergent connectivity patterns between awake and anesthetized mice, across imaging modalities and timepoints, **Table 1**.

**Table 1.**
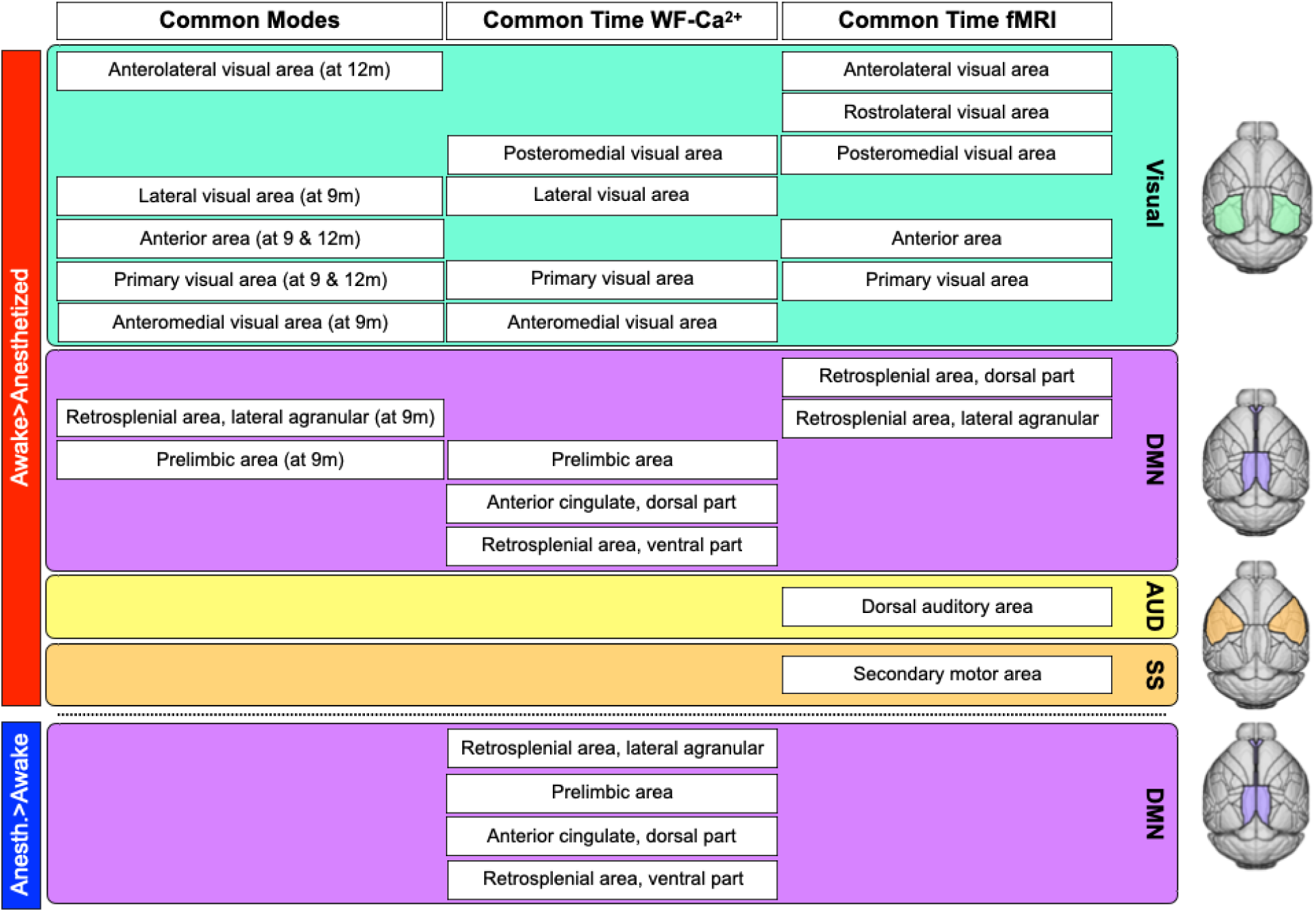
Nodes showing connectivity differences in awake vs. anesthetized mice in multimodal data.

Nodes that showed greater connectivity in awake relative to anesthetized mice at 9m in both imaging modalities included visual areas and regions within the default mode network (DMN), **Table 1** (column 1). At the 12m imaging timepoint, a subset of visual areas showed the same pattern. In the WF-Ca^2+^ imaging data, greater connectivity in awake relative to anesthetized mice was recovered at both imaging timepoints in visual areas and regions within the DMN, **Table 1** (column 2). The opposite pattern, increased connectivity in anesthetized relative to awake mice, was also observed with WF-Ca^2+^ imaging within the DMN. In parallel, the fMRI data showed greater connectivity in awake relative to anesthetized mice at both timepoints in visual areas and regions within the DMN as well as in the dorsal auditory (AUD) area, and secondary motor area, **Table 1** (column 3).

Overall, there were several brain regions that showed consistent changes in connectivity between awake and anesthetized mice across imaging modalities and timepoints. These were dominated by increased connectivity in visual areas and regions within the DMN. However, there were also clear instances when the two imaging modalities showed different or diverging patterns. As above (**2.4.1**), these observations were well replicated when the 6m (anesthetized) imaging timepoint was used in-place of the 4m imaging timepoint.

#### 3.4.3 A priori cortical brain networks showed differences in connectivity between awake and anesthetized mice that differed between imaging modalities

Brain regions were organized into networks, **Fig. S3**. In the previous subsection (**3.4.2**) brain regions showing differences in connectivity between awake and anesthetized mice were identified alongside the networks they belonged to (bottom-up). When *a priori* networks, or inter-network pairs, were considered (top-down) in a linear mixed-effects model (see **2.8.3**) with *State* (awake, anesthetized), *Hemisphere* (intra-vs. inter-hemisphere), and *Network* as fixed effects and *Mouse* as a random effect, differences in connectivity between awake and anesthetized mice within or between networks revealed more divergent patterns between imaging modalities, **Fig. 4B&C**.

From WF-Ca^2+^ imaging, data from the 9m (awake) imaging timepoint, relative to the 4m (anesthetized) imaging timepoint, showed a significant main effect for network (F_3,264_=29.14, p=2.6414e-16), as well as a significant interaction between network x hemisphere (F_3,264_=3.57, p=0.015) and state x hemisphere (F_1,264_=15.62, p=9.9326e-05). Similarly, data from the 12m (awake) imaging timepoint, relative to the 4m (anesthetized) imaging timepoint, showed a significant main effect for network (F_3,288_=34.99, p=2.6152e-19), state (F_1,288_=5.36, p=0.02) as well as a significant interactions between network x hemisphere (F_3,288_=4.29, p=0.006), state x hemisphere (F_1,288_=26.71, p=4.4275e-07), and hemisphere x network x state (F_3,288_=4.12, p=0.007), indicating that the effect of wakefulness on functional connectivity depended on both the network and whether connectivity was measured within or across hemispheres.

Overall, differences in *a priori* network connectivity between awake and anesthetized mice were more prominent inter-hemisphere than intra-hemisphere in the WF-Ca^2+^ imaging data, **Fig. 4C**. Motor, somatosensory and auditory networks showed decreases in inter-hemispheric connectivity at the 9 and 12m awake imaging timepoints, relative to the 4m anesthetized imaging timepoint (p<0.05, pairwise post-hoc test, Benjamini-Hochberg); the visual network showed decreases at 12m compared to 4m (p<0.001). The DMN showed the same patten at the 12m imaging timepoint intra-as well as inter-hemisphere, whilst no other differences in intra-hemispheric network connectivity were observed with WF-Ca^2+^ imaging. Permutation testing confirmed that the observed awake - anesthetized differences exceeded those expected by chance for inter-hemispheric connectivity in Motor, Somatosensory/Auditory, and DMN (the latter only for 12m vs. 4m), as well as intra-hemispheric connectivity in DMN (at 12m vs. 4m, p<0.05), whereas the remaining comparisons were not significant (**Fig. S11A**, left).

Dissimilarly, fMRI showed widespread increases in *a priori* network intra-hemispheric as well as inter-hemispheric connectivity at the 9 and 12m imaging timepoints, relative to the 4m imaging timepoint. As above, the fMRI data acquired at 4 and 9m showed a significant main effect of network (F_3,264_=14.15, p=1.4093e-08), as well as a significant effect of hemisphere (F_1,264_=87.13, p=4.335e-18) as well as state (F_1,264_=66.47, p=1.4429e-14). Significant interactions were observed between network x hemisphere (F_3,264_=5.47, p=0.0011681), state x network (F_3,264_=16.57, p=6.8875e-10), and hemisphere x network x state (F_3,264_=16.99, p=4.0364e-10).

Very similar results were found using the fMRI data acquired at the 4 and 12m imaging timepoints. Significant main effects of network (F_3,288_=18.04, p=9.339e-11), hemisphere (F_1,288_=111.13, p=3.4448e-22) as well as state (F_1,288_=53.64, p=2.4163e-12) were observed. Further, significant interaction between network x hemisphere (F_3,288_=6.97, p=0.00015271), state x network (F_3,288_=10.73, p=1.0451e-06), and hemisphere x network x state (F_3,288_=3.67, p=0.012761) were found. Of note, fMRI inter-hemispheric connectivity in *a priori* networks showed a divergent pattern, with connectivity in motor areas being increased at the 9 and 12m imaging timepoints (p<0.01 for both), and somatosensory/auditory showing decreased connectivity, relative to the 4m imaging timepoint (p<0.01 only for 9m, 12 m not surviving). The robustness of these observations is supported by permutation tests where subject and condition (awake or anesthetized states) were randomly shuffled and compared to the true observed values (n=1,000 iterations), across all networks, **Fig. S11A**, right.

Together, these findings suggest that the transition to wakefulness induces broad reorganization of large-scale BOLD functional networks, while WF-Ca^2+^ connectivity is modulated in a more network-specific manner, predominantly through reduced inter-hemispheric synchrony.

#### 3.4.4 Node and network analyses can lead to seemingly different conclusions

When brain regions (bottom-up) were considered (**3.4.2**), the nodes identified as playing a prominent role in distinguishing awake from anesthetized mice included visual areas as well as regions within the DMN, **Table 1**. Although some differences in the corresponding networks were recovered (**3.4.3**) – intra/inter-hemispheric DMN connectivity at the 12m imaging timepoint, in the WF-Ca^2+^ imaging data, and intra-hemispheric visual connectivity at the 9 and 12m imaging timepoints in the fMRI data – they were not among the more salient effects. This is a result of considering a network in its entirety, which can obscure contributions from individual nodes, opposing contributions from different nodes, or connections, within the same network (e.g., DMN, **Table 1**), and contributions from inter-network connectivity, **Fig. 4B**.

#### 3.4.5 Whole-brain a priori networks observed with fMRI show widespread differences between awake and anesthetized mice that are independent of mouse age

Besides the analyses at the cortical level highlighted above, seven *a priori* networks which span the whole-brain were evaluated using the fMRI data, **Fig. S3**. As in the multimodal analyses of cortical connectivity, widespread differences in whole-brain connectivity between awake and anesthetized mice were observed at the 9 and 12m imaging timepoints relative to the 4m (anesthetized) imaging timepoint, **Fig. S8**. Similarly, an overall decrease in these differences was apparent in the data collected at the 12 relative to 9m imaging timepoint, **Fig. S8A**. When *a priori* network connectivity, both intra- and inter-hemisphere, was compared across imaging timepoints, **Fig. S8B**, both patterns of increased connectivity in awake relative to anesthetized mice and the opposite pattern were observed. For the 9 versus 4m imaging timepoint comparison, significant main effects of hemisphere (F_1,462_=15.36, p=0.0001), network (F_6,462_=20.18, p=5.3526e-21), and state (F_1,462_=67.17, p=2.4706e-15) were observed, alongside significant interactions between state x hemisphere (F_1,462_=71.19, p=4.1987e-16), state x network (F_6,462_=23.48, p=3.0285e-24), hemisphere x network (F_6,462_=3.78, p=0.001) and network x hemisphere x state (F_6,462_=22.12, p=6.4512e-23).

Similarly, for the 4 versus 12m imaging timepoint comparison significant main effects of hemisphere (F_1,504_=22.00, p=3.5142e-06), network (F_6,504_=28.90, p=9.3029e-30), and state (F_1,504_=35.6, p=4.5659e-09) were observed, alongside significant interactions between state x hemisphere (F_1,504_=45.85, p=3.5613e-11), state x network (F_6,504_=14.95, p=9.1364e-16), hemisphere x network (F_6,504_=5.41, p=1.9386e-05) and network x hemisphere x state (F_6,504_=7.81, p=4.712e-08).

Overall, intra-hemisphere, connectivity in awake relative to anesthetized mice increased, with the exception of within the DMN, which showed no effect, and the striatal network which showed increased connectivity in anesthetized relative to awake mice. Conversely, inter-hemispheric connectivity was more widely increased in anesthetized relative to awake mice, besides thalamic and basal ganglia networks.

Taken together, extending the analysis to whole-brain networks revealed a similar pattern to cortex-only results, with wakefulness driving widespread, network-dependent reorganization of BOLD functional connectivity that was largely preserved across awake sessions. As above, the robustness of these observations was supported by permutation tests where subjects and condition (awake or anesthetized) were randomly permuted, **Fig. S11B**. Furthermore, when the 6m (anesthetized) imaging timepoint was used, in-place of 4m, similar results were obtained.

The properties of the dataset (**2.5**) also allowed for a linear mixed model to be generated using fMRI, but not WF-Ca^2+^ imaging data, which jointly accounted for differences in connectivity associated with animal condition (awake or anesthetized) and age. Two models were generated. One using data from the cortex, matching the WF-Ca^2+^ imaging FOV (**Fig. S10A**), and one using data from the whole-brain, **Fig. S10B**. Both showed that the effects of age on differences in connectivity were minimal compared to the effects of condition (awake versus anesthetized). We also observed a significant interaction between age and state, which indicated that the effect of state on functional connectivity was not constant across imaging sessions. Specifically, state-related differences were more pronounced at the first relative to second awake imaging session (**Fig. S10**, right).

### 3.5 Cross-modality correlation of WF-Ca^2+^ and fMRI connectivity

Diverging patterns in intra- and inter-hemispheric connectivity across cortical networks in WF-Ca^2+^ and fMRI data (**3.5.3**) motivated an examination of cross-modality connectivity agreement within and between networks, **Fig. 5A**, as well as intra- and inter-hemisphere, **Fig. 5B**. WF-Ca^2+^ and fMRI connectomes were separated into cortical networks within right and left hemispheres and vectorized. Due to its small size (two regions per hemisphere), the motor network was excluded. Cross-modality correlations (Pearson’s R) were computed for each pair of networks (and hemispheres) at each imaging timepoint.

**Fig. 5.**
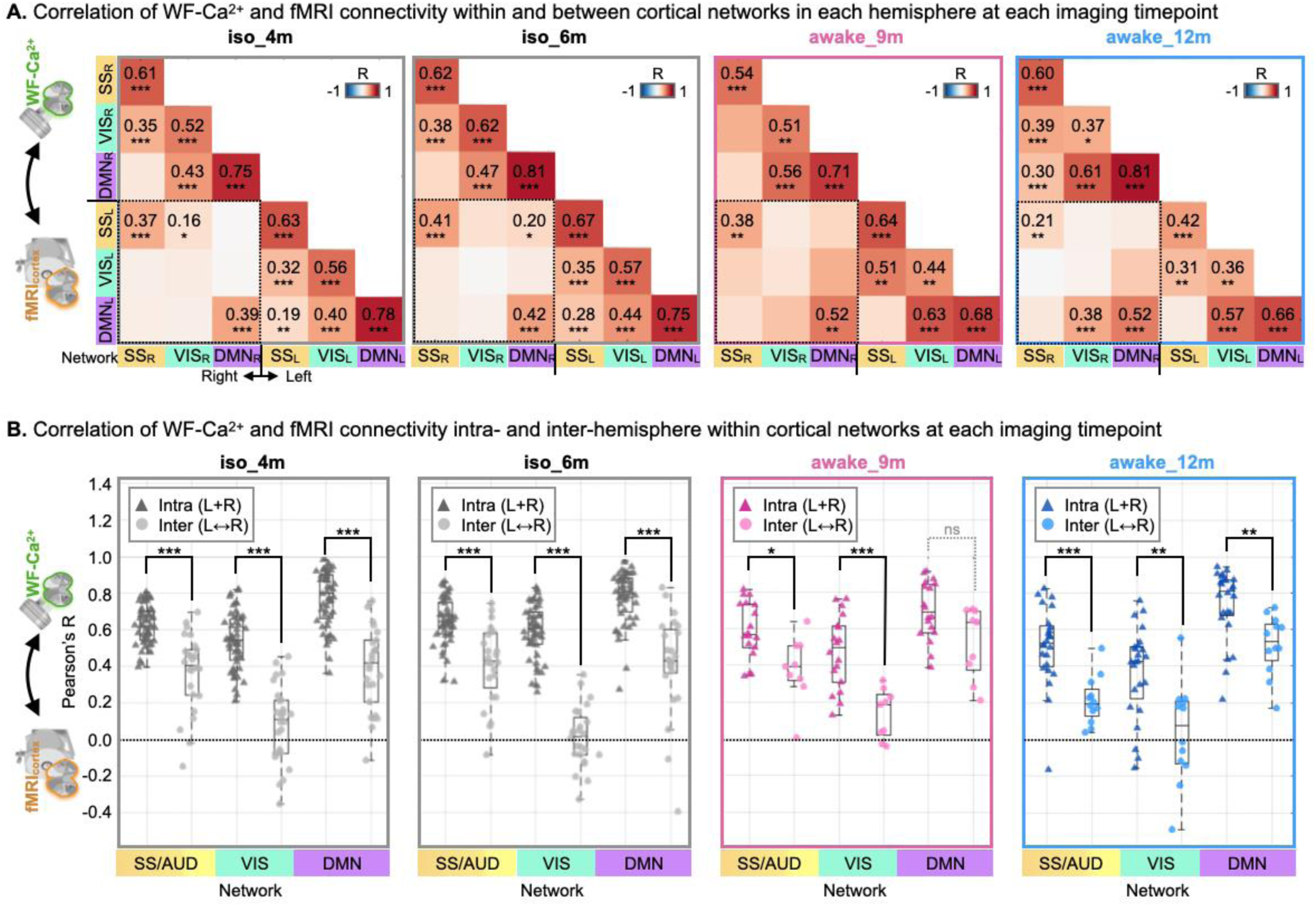
Correlation of WF-Ca^2+^ and fMRI connectivity across cortical networks and imaging timepoints. **A.** At each imaging timepoint (4, 6, 9, and 12m), matrices showing the mean correlation of WF-Ca^2+^ and fMRI connectivity within and between cortical networks. Data from right and left hemispheres, as well as inter-hemisphere (black dotted lines), are shown. **B.** Scatterplots showing intra- and inter-hemisphere cross-modality correlation of WF-Ca^2+^ and fMRI connectivity within cortical networks (intra/inter are denoted by triangles/circles, respectively) at each imaging timepoint. Spatial resolution: raw WF-Ca^2+^ = 25×25 μm^2^; raw fMRI = 310×310×310 μm^3^; Temporal resolution: raw WF-Ca^2+^ = 10Hz; raw fMRI = 0.55Hz. Corrected p-values are displayed: *p<0.05, **p< 0.01, ***p<0.001.

#### 3.5.1 Patterns in cross-modality connectivity were well-preserved across imaging timepoints and between conditions (awake vs. anesthetized states)

At both anesthetized, 4 and 6m, and awake, 9 and 12m, imaging timepoints, remarkably similar inter-modality connectivity patterns were observed. Within cortical networks, in both hemispheres, high inter-modality correlations were recovered **Fig. 5A** (values along the diagonal) – especially within the DMN. Across hemispheres, there was much less inter-modality agreement, **Fig. 5A** (dashed line box).

#### 3.5.2 Across cortical networks, and imaging timepoints, correlation strength between WF-Ca^2+^ and fMRI connectivity was lower inter-versus intra-hemisphere

Using connectomes generated from each run (10mins), correlation strength between WF-Ca^2+^ and fMRI connectivity for each cortical network, intra- and inter-hemisphere were plotted, **Fig. 5B**. Across cortical networks and imaging timepoints, intra-hemisphere connectivity across imaging modalities was more correlated than inter-hemisphere connectivity, with the sole exception of the DMN at the 9m imaging timepoint. This pattern was reminiscent of observations made in our previous work, which considered node-connectivity across imaging modalities intra- and inter-hemisphere (Lake *et al*., 2020).

## 4. Discussion

This work establishes a biphasic acclimation protocol for longitudinal simultaneous WF-Ca^2+^ and fMRI data collection in awake adult mice. The experimental framework incorporates elements from a recent systematic review where we highlighted the importance of comparing data collected from awake and anesthetized animals alongside a critical evaluation of signal quality given the paucity of such investigations in the current literature (Mandino *et al*., 2024). Beyond the technical advancement needed for performing these experiments, this work provides a novel characterization of large-scale brain connectivity differences between awake and anesthetized mice using unique multimodal data that are specifically well-suited to network-based analyses given their large and overlapping coverage of the brain. Coincident inter-modal convergent and divergent patterns in large-scale functional organization are revealed and considered within the context of previously observed differences in data collected from awake and anesthetized mice which used either WF-Ca^2+^ (Brier *et al*., 2019; Wright *et al*., 2017) or fMRI (Dinh *et al*., 2021; Fadel *et al*., 2022; Gutierrez-Barragan *et al*., 2022; Tsurugizawa *et al*., 2021).

Each element of the biphasic acclimation protocol was based on evidence gleaned through our systematic review (Mandino *et al*., 2024). This included the 14-day duration of Phase-one: ‘Initial Progressing Training’, gradual and incremental introduction of stressors, use of a food-reward, biphasic design, use and key elements of a mock scanner environment (e.g., recorded scanner noise), as well as use of the real system for acclimation purposes. These efforts contributed to achieving motion estimates (i.e., |FD|) which were comparable to previous high-quality work at both timepoints where mice were imaged whilst awake. A detailed comparison in **Fig. S4**, adapted from our recent systematic review (Mandino et al., 2024), places these measurements in the context of the wider literature. Overall, the proportion of frames exceeding our motion threshold (∼22% in the first awake imaging session) fall within the reported range. This estimate decreases (to ∼10%) following the second phase of acclimation. The decrease in subject motion which followed Phase-two: ‘Refresher Training’, a full two-months following the initial phase, **Fig. 2C**, could motivate an investigation into a more distributed, rather than intensive, acclimation protocol as part of future work. That the study design implemented here did not include a group which underwent Phase-one: ‘Initial Progressive Training’ prior to the 12m imaging timepoint precludes concluding that a biphasic protocol reduces subject motion based on the current study results alone.

Simultaneous WF-Ca^2+^ imaging showed minimal motion in the X–Z plane (the only estimate possible using these 2D data). This is in-line with the observation that the majority of motion in the fMRI data is along the Y-axis, which is “invisible” to WF-Ca^2+^ imaging, **Fig. S2**. Movements observed along shared spatial dimensions (e.g., Z) which are recovered by fMRI motion correction only likely arise from body movements that manifest as ‘head motion’ in the fMRI signal and have no correlate in the WF-Ca^2+^ imaging data. These are caused by induced B_0_ inhomogeneities. A deeper phenotyping of how subject motion manifests in multimodal data and its impact on measures of brain function will be the focus of future work. Other avenues include the future use of a body restraint system (Laxer *et al*., 2025; Mandino *et al*., 2024; Russo *et al*., 2021) (as the mouse body was unrestricted in this study), magnetic field correction (Zeng *et al*., 2022), or the implementation of novel MRI-acquisition strategies that are insensitive to body movement (Gozzi *et al*., 2025; Mangia *et al*., 2025; Paasonen *et al*., 2020).

In the fMRI data, tSNR at all timepoints was comparable to estimates from anesthetized animals in a recent multi-center study (see **Fig. 1** in (Grandjean *et al*., 2020)). Still, the tSNR in the fMRI data acquired from awake mice was lower than in data acquired from anesthetized mice. In the corresponding WF-Ca^2+^ imaging data, tSNR showed the opposite pattern, **Fig. 2D**. This is unsurprising given the distinct signal and noise sources, including head and body motion (**3.1.1**), which affect tSNR differentially in each modality (Keilholz *et al*., 2017; Lecoq *et al*., 2021; Zhang *et al*., 2023).

Acclimation protocols for undergoing imaging whilst awake aim to reduce animal motion, improve tSNR, and reduce animal stress through exposure to the conditions of the experiment without any adverse events. Ideally, through acclimation, the animal learns that the procedures are not harmful, and this reduces stress. However, over-training has been posited as equally undesirable as it can result in a depressive phenotype due to prolonged, low-level, stress (Herrera-Portillo *et al*., 2025). Striking the ‘ideal’ balance between these extremes–both of which have known effects on brain function (Akil *et al*., 2023; Herrera-Portillo *et al*., 2025; Lupinsky *et al*., 2025) is a challenging and currently unsolved problem. Stress, both acute and chronic, is difficult to measure in mice and has not been thoroughly characterized in the existing mouse fMRI literature (Mandino *et al*., 2024). Further, these concerns have been typically ignored in WF-Ca^2+^ imaging experiments due to the environment being considered much less stressful overall, with a few notable exceptions (Juczewski *et al*., 2020; Nietz *et al*., 2022; Rynes *et al*., 2021). A thorough investigation of the effects and presence of acute and chronic stress was beyond the scope of this investigation. However, animal body weight, measured daily during acclimation and imaging, was used as a proxy for long-term stress (Kuti *et al*., 2022). Every animal in this study maintained a stable weight throughout acclimation and imaging, with no differences between awake and anesthetized groups, **Fig. S1**. Future work will include more comprehensive measures of acute and chronic stress using behavioral monitoring, heart rate variability (Tsurugizawa *et al*., 2020; Yoshida *et al*., 2016), and/or blood corticosterone measurements (Gutierrez-Barragan *et al*., 2022; Mandino *et al*., 2024). Overall, the awake mouse fMRI literature has always included some amount of habituation. Most acclimation protocols span 10–14 days while shorter durations are associated with increased motion and data loss (Mandino et al., 2024). In designing our study, we elected to operate within this established range to preserve data quality. Ongoing efforts across laboratories will be important for further refinement of generalizable acclimation strategies and a deeper understanding of the impacts different approaches have on data quality and experimental efficiency.

So-called “specific” connectivity was recently introduced by Grandjean and colleagues ((Grandjean *et al*., 2020; Grandjean *et al*., 2023)) as a metric for assessing rodent fMRI data quality. It is based on quantifying two “biologically plausible” measures of functional connectivity, one posited to show high connectivity (SSp_R_ ↔ SSp_L_), and the second posited to show low, absent, or negative connectivity (SSp_R_ ↔ VIS_R_), based on the physical/structural connectedness of these brain regions, **Fig. 3A**. A large fraction of the fMRI data collected from anesthetized mice in this study was categorized as “high-quality” based on this metric, **Fig. 3C**. Functional MRI data acquired from awake mice, encouragingly, showed similarly high rates despite lower tSNR. Notably, the specificity results across all fMRI data were aligned with previous literature (∼80% in a recent multi-center mouse fMRI study (Grandjean *et al*., 2020)).

Extending this assessment to the simultaneously acquired WF-Ca^2+^ imaging data collected from anesthetized mice, showed similar results, with specific connectivity across almost the entire dataset, both in the anesthetized and awake states (**Fig. 3B**). Given that this metric is based on a viral tracer map, it should be more closely related to WF-Ca^2+^ imaging results, than those gleaned from fMRI data (despite this metric having been originally developed for assessing fMRI data quality). Overall, this weighs-in on a long-standing debate regarding the relatedness of brain structure and function in both the human (Baum *et al*., 2020; Fotiadis *et al*., 2024; Popp *et al*., 2025) and mouse (Benozzo *et al*., 2024; Grandjean *et al*., 2017; Melozzi *et al*., 2019) literature.

Data from awake, relative to anesthetized mice, showed widespread increases in intra-hemispheric connectivity strength alongside decreases in inter-hemispheric connectivity strength within *a priori* cortical and whole-brain networks, **Fig. 4C**, **S8 & S10**, with a few noted exceptions (**3.4**). Similar patterns have been previously reported using both mice (Dvorakova *et al*., 2024; Tsurugizawa *et al*., 2020) and rats (Paasonen *et al*., 2018), and are purported to reflect enhanced network segregation and reduced global synchrony with wakefulness (Desai *et al*., 2011; Fadel *et al*., 2022; Gutierrez-Barragan *et al*., 2022; Jonckers *et al*., 2014; Tsurugizawa *et al*., 2021; Yoshida *et al*., 2016), see **Table 3** in (Mandino *et al*., 2024). Although these patterns were recovered at both the 9 and 12m imaging timepoints, the effects were stronger, in the fMRI data, at the 9m imaging timepoint. Using the fMRI data, it was inferred that the attenuation of these effects were not attributable to animal age, **Fig. S10**. The presence of a significant age × state interaction further suggests that the impact of wakefulness on large-scale network organization is dynamic, rather than fixed, potentially reflecting ongoing adaptation to the imaging environment. Although, it is possible that the completion of additional acclimation was responsible, this will need to be validated as part of future work. It is also worth noting that the effects of age on connectivity observed in the present work are milder, albeit following the same trend, than what we have reported previously (Mandino *et al*., 2025b), despite overlap between datasets (**Methods**). The likely difference being the application of global signal regression in our current analysis and its omission in prior work (compare **Fig. S8B** with **Fig. 2** in (Mandino *et al*., 2025b)). This may indicate that age-related effects are present within the global signal. Future work will aim to examine this possibility.

Relative to observations made using fMRI, simultaneously acquired WF-Ca^2+^ imaging showed more stable intra-hemisphere connectivity strength between awake and anesthetized mice within *a priori* cortical networks, while differences in inter-hemispheric connectivity strength showed a mixture of convergent (motor and visual) as well as divergent (motor and DMN) patterns, **Fig. 4C**. It was also apparent, that the WF-Ca^2+^ imaging data showed more widespread differences in connectivity strength between awake and anesthetized mice at the 12m imaging timepoint (whereas fMRI showed the opposite pattern), **Fig. 4A**. However, closer inspection of the matrices in **Fig. 4A** derived from the WF-Ca^2+^ imaging data, especially at the 12m imaging timepoint, revealed that averaging within *a priori* networks likely obscured a range of differences in connectivity strength between awake and anesthetized mice. Most networks contained nodes showing both increased and decreased connectivity strength between awake and anesthetized mice which canceled when averaged within network. There were also noticeable differences between awake and anesthetized mice in inter-network connectivity. Some of these features were captured when differences in brain region connectivity was quantified (**3.4.2**) but further investigation is warranted. In this study, both *a priori* networks, and brain regions (derived from the Allen Atlas (Atlas, 2004; Wang *et al*., 2020)), were adopted for the purpose of comparing to the existing literature. In future work, a less constrained approach will be implemented, as we have done previously (Vafaii *et al*., 2024), which will support a more data driven characterization of these features.

The correlation of WF-Ca^2+^ and fMRI connectivity within and between *a priori* networks, separating inter- and intra-hemispheric connections (**Fig. 5**), aligns well with our earlier work (see **Fig. 6B** in (Lake *et al*., 2020)). High inter-modality correlations are recovered within-network in both hemispheres (diagonal, **Fig. 5A**) alongside lower or absent correlations between networks and across hemispheres (off-diagonal, **Fig. 5A**). In our previous work (Lake *et al*., 2020), mice were imaged under anesthesia which was suspected as the reason for the lack of inter-hemispheric connectivity agreement between WF-Ca^2+^ and fMRI. In this study, this hypothesis was disproven. Across cortical networks, and imaging timepoints, the same pattern was reproduced regardless of whether mice were anesthetized or awake during imaging, **Fig. 5B**. Further examination of this persistent effect will aim to quantify the role that connection length plays in determining inter-modality connectivity agreement. We anticipate that this analysis will benefit from using a brain atlas comprised of smaller and more similarly sized brain regions.

In sum, this work introduces longitudinal, simultaneous WF-Ca^2+^ and fMRI in awake mice. The biphasic acclimation protocol yields high-quality multimodal data appropriate for quantifying cortex-wide and whole-brain (fMRI) activity patterns with complementary sources of contrast. Novel aspects of patterned brain activity in awake and anesthetized mice were captured. Convergent and divergent activity patterns were quantified across spatial scales and contrasted between imaging timepoints and modalities. This work sets the stage for future studies in awake mice, where a wider spectrum of behaviors and brain states can be examined in healthy animals and models of disease. Finally, the present work focuses on connectivity- and network-level analyses to characterize large-scale organization across modalities and brain states, providing a foundation for cross-modal and cross-state comparisons. Building on this framework, future work will incorporate time-resolved analyses of both WF-Ca^2+^ and fMRI signals, including spontaneous activity patterns, temporal coupling, and state-dependent dynamics across modalities. These extensions will further enrich the multimodal framework and offer new opportunities to disentangle the relationship between neuronal and vascular signals in awake conditions.

## Acknowledgments

Drs. Joel Greenwood, Paul Shamble, and Omer Mano from Neurotechnology Core and the team members of the Magnetic Resonance Research Center at Yale University for their technological expertise. Dr. Joanes Grandjean for critical discussion. Dr. Yonghyun Ha for hardware support. Dr. Ali S. Hamodi and Dr. Michael C. Crair for experimental expertise.

## **5.** Funding

CH was supported by NIH R25MH119043. EMRL, FM, SMS were supported by NIH R01 NS130069-01.

## **6.** Author contributions

Conceptualization: FM, SMS, EMRL.

Methodology: FM, XS, TK, CH, EMRL.

Investigation: FM, EMRL.

Visualization: FM, EMRL.

Funding acquisition: SMS, EMRL.

Project administration: FM, EMRL.

Supervision: EMRL.

Writing—original draft: FM.

Writing—review & editing: all co-authors.

## **7.** Competing interests

XP owns small stakes in Veridat and Tytonix which are tech startup companies.

## **8.** Data and materials availability

All data and codes used in this manuscript will be rendered available by the corresponding authors upon request.

## Notes

### Summary of Updates

This version implements reviewers' comments including additional methodology explaination, and analyses. Statistical analyses have been carried out to include linear-mixed models to account for aging effects across analyses. Further WF-Ca2+ analyses using the higher frequencies (Calcium 'fast') are now also included.

