## Supplementary for "Biphasic acclimation for simultaneous wide-field fluorescent Ca^2+^ imaging and fMRI of awake mice"

**Supplementary material**

**Supplementary Tables**

**Table S1.** Animal cohort breakdown, all data used in analyses

| | **Age** (m) | **Longitudinal *** (N = animal, n = runs) | **Cross-sectional** | **State** |
|---|---|---|---|---|
| **iso_4m** | 4m | N = 5/4+2** m/f, n = 27 | N.A. | Anesthesia |
| **iso_6m** | 6m | N = 5/4 m/f, n = 27 | N.A. | Anesthesia |
| **iso_9m** | 9m | N = 0/1ǂ+1** m/f, n = 3 | N = 4/2 m/f, n = 18 | Anesthesia |
| **awake_9m** | 9m | N = 3/2 m/f, n = 15 | N.A. | Awake post progressive training |
| **iso_12m** | 12m | N = 0/1ǂ+1** m/f, n = 3 | N = 2/2 m/f, n = 12 | Anesthesia |
| **awake_12m** | 12m | N = 2/2 m/f, n = 12 | N.A. | Awake post refresher training |

*Also have WF-$Ca^{2+}$ data
ǂ mouse (and cohort) excluded from analyses (because single animal in that cohort)
**mouse excluded during analyses due to anatomical issues
m = months

**Table S2.** Censoring of functional data (fMRI)

| Session | Mean Retained Frames | SD Retained Frames | Mean Censored Frames | SD Censored Frames |
|---|---|---|---|---|
| **iso_4m** | 319.42 | 7.7881 | 15.577 | 7.7881 |
| **iso_6m** | 319.75 | 6.2433 | 15.25 | 6.2433 |
| **awake_9m** | 214.89 | 38.218 | 120.11 | 38.218 |
| **iso_9m** | 316.14 | 9.1394 | 18.857 | 9.1394 |
| **awake_12m** | 244.33 | 37.257 | 90.667 | 37.257 |
| **iso_12m** | 312.33 | 12.01 | 22.667 | 12.01 |

### 2 Supplementary Figures

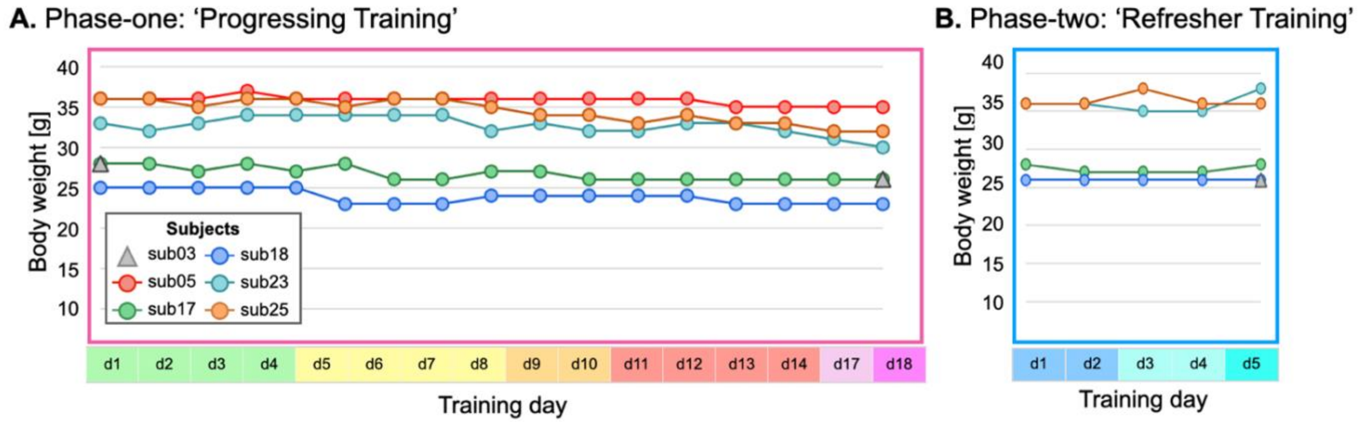


**Fig. S1. Mouse body weight during biphasic acclimation training. A.** Body weight measured daily during Phase one: ‘Progressive training’ for acclimating naïve mice to undergoing multimodal imaging whilst awake. A timeline for Phase one acclimation training is shown in **Fig. 1E**. Mice being acclimated are plotted in color. Body weight from a control mouse that did not undergo acclimation training is plotted in grey. **B.** As in (**A.**) for Phase two: ‘Refresher training’ for re-acclimating mice to undergoing multimodal imaging whilst awake. A timeline for Phase two acclimation training is shown in **Fig. 1G**.

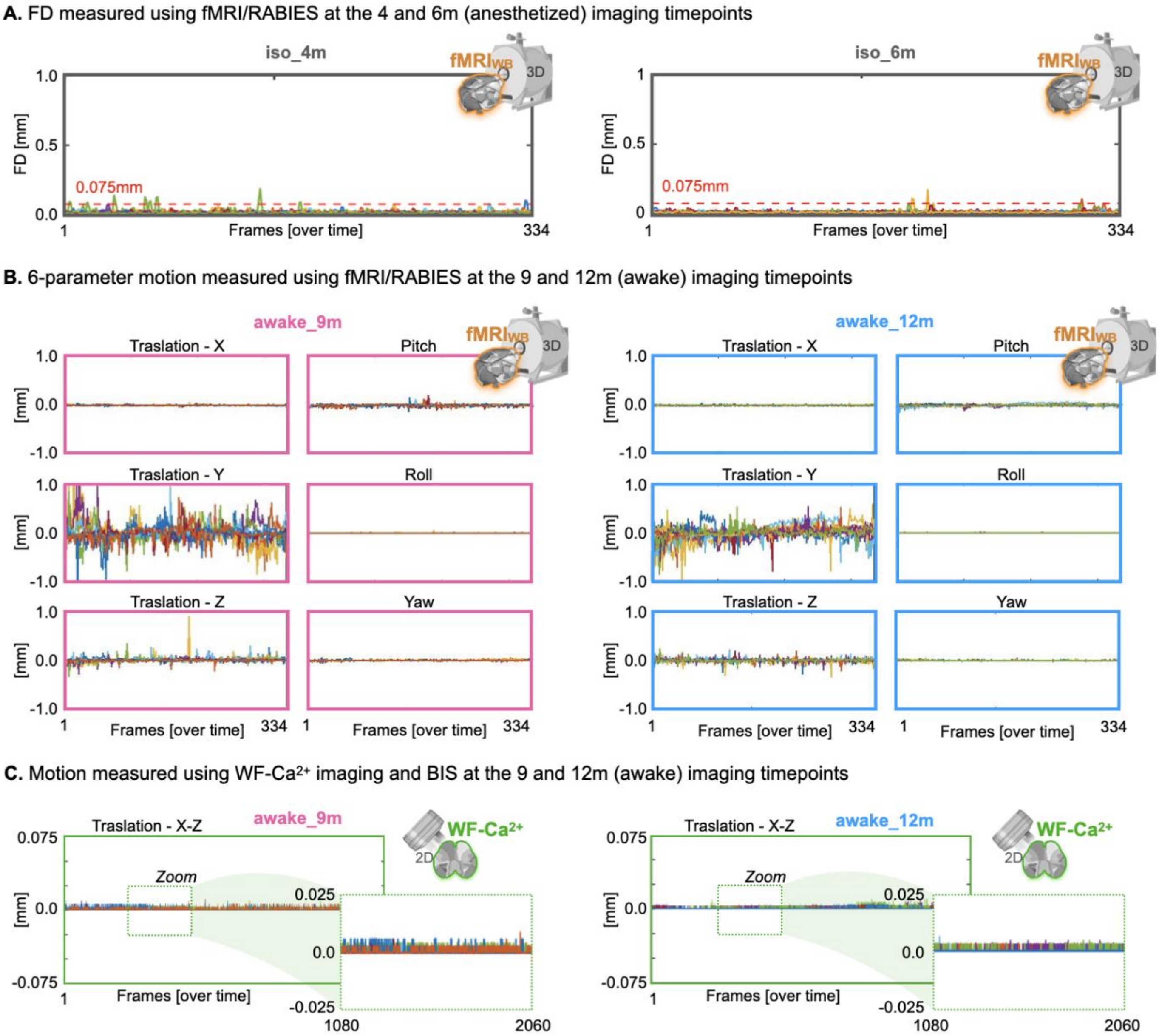


**Fig. S2. Motion in multimodal imaging data. A.** FD measured using fMRI data and RABIES the at 4 and 6m (anesthetized) imaging timepoints. Corresponding FD measures for the 9 and 12m anesthetized and awake imaging timepoints are shown in **Fig. 2B**. **B.** Motion from the 9 and 12m (awake) imaging timepoints broken down into 6-parameters. **C.** Motion in the WF-$Ca^{2+}$ imaging data, estimated using BIS software, at the 9 and 12m (awake) imaging timepoints.

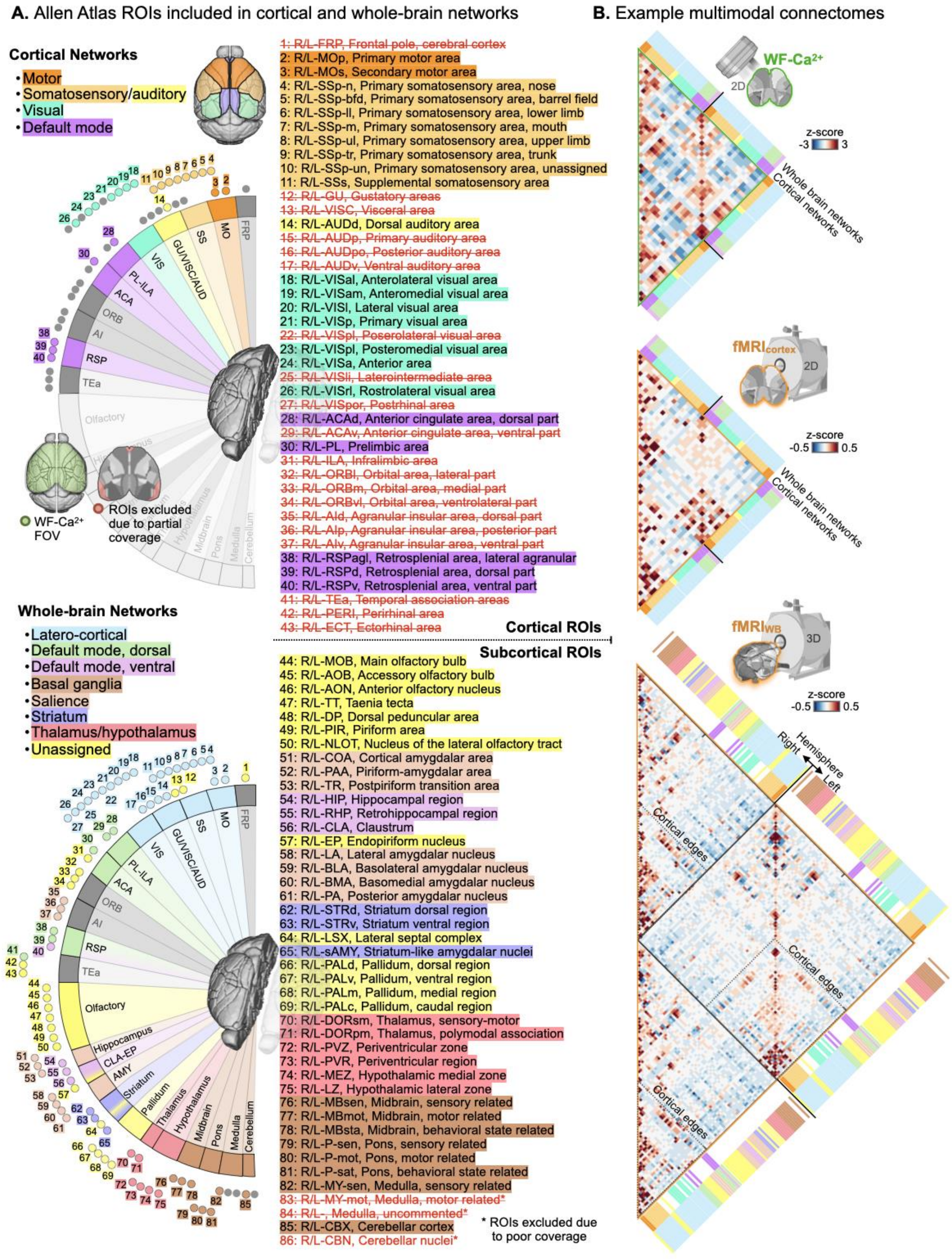


**Fig. S3. Allen Atlas derived brain regions and a priori cortical and whole-brain networks. A. (top)** Brain regions visible in the WF-Ca$^{2+}$ imaging FOV used in all cortical network analyses. Regions were subdivided into four cortical networks. **(bottom)** Brain regions visible in fMRI data used in all whole-brain network analyses. Regions were subdivided into seven networks. **B.** Averaged connectomes from multimodal data including cortical (top) and whole-brain examples. Network assignments are color-coded.

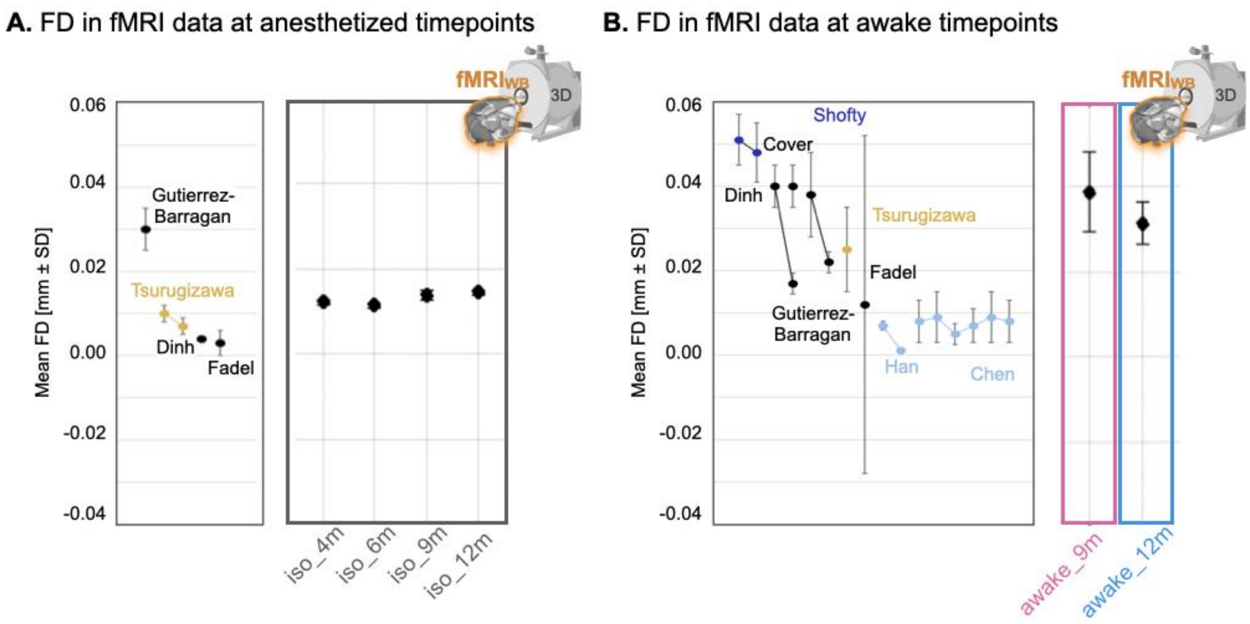


**Fig. S4. FD in fMRI data estimated with RABIES compared to current literature. A,B:** Left panels show reproduced values from **Figure S8** in a systematic review of awake mouse fMRI (Mandino *et al.*, 2024). Right panels show data from the current study. **A.** Mean FD in fMRI data acquired from anesthetized mice. **B.** Mean FD in fMRI data acquired from awake mice.

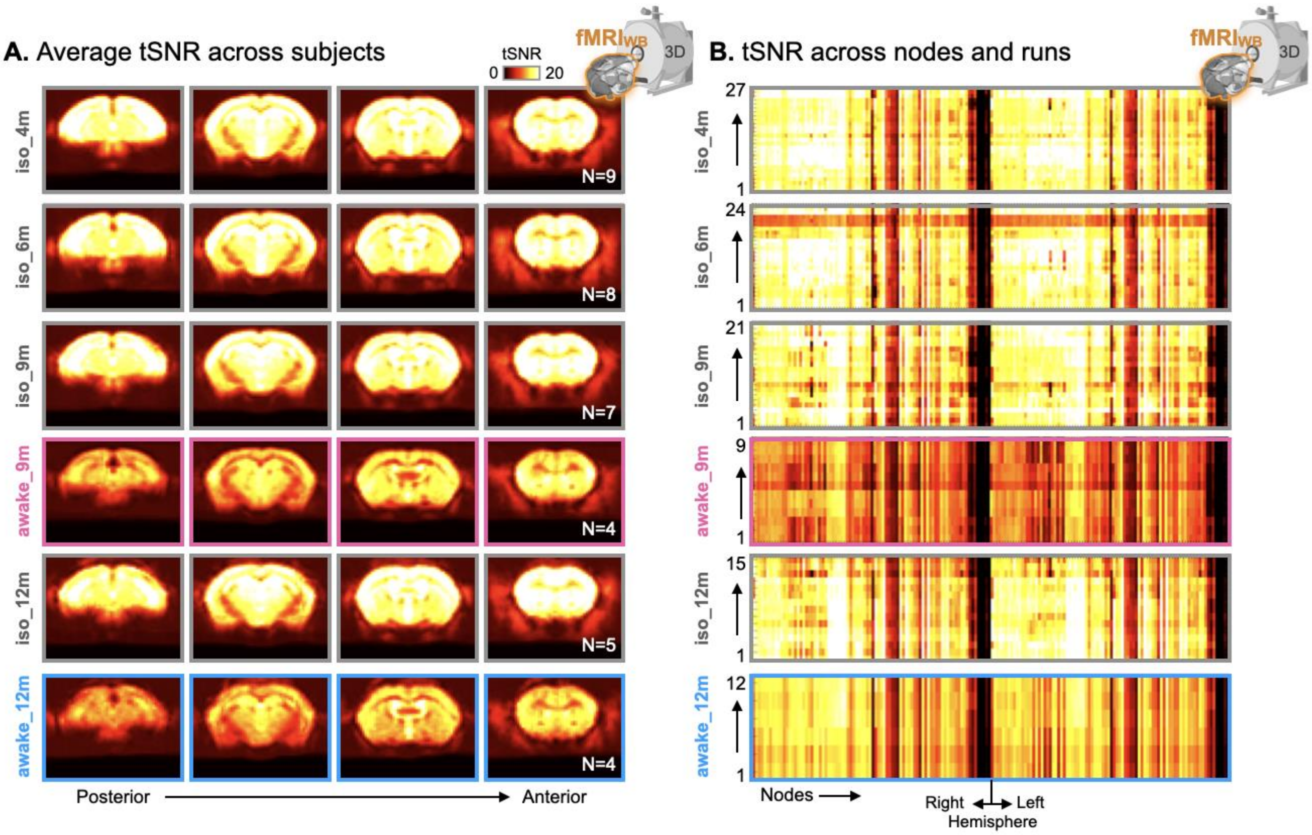


**Fig. S5. Temporal SNR of fMRI data measured with RABIES. A.** Example group averaged tSNR maps estimated with fMRI/RABIES at each imaging timepoint. **B.** Carpet plots output by RABIES which show tSNR estimated for each brain region (node), and run, at each imaging timepoint.

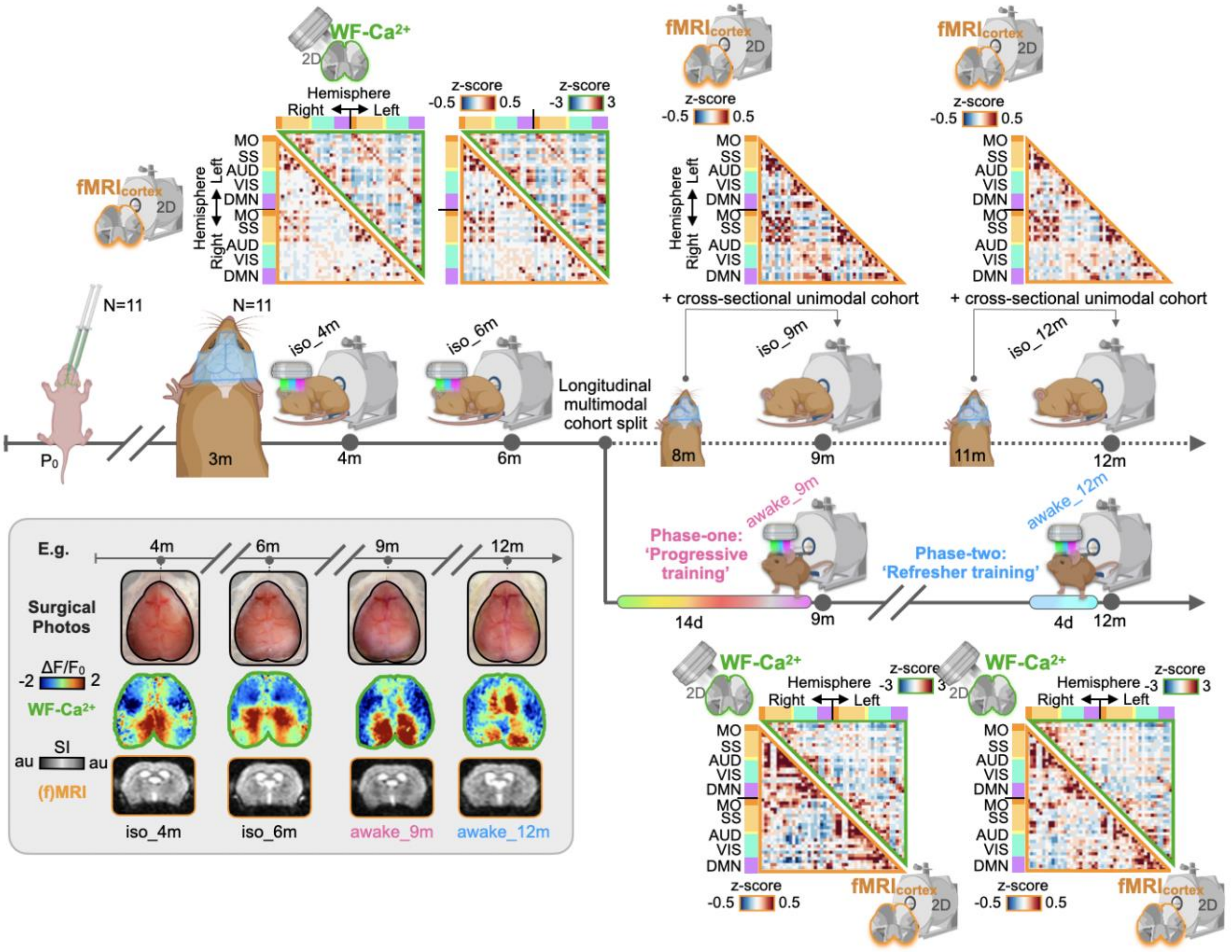


**Fig. S6. Experimental timeline with example data.** The timeline is copied from **Fig. 1A.** Above and below each multimodal imaging timepoint, averaged connectomes using WF-$Ca^{2+}$ (green outline) or fMRI (orange outline) data from the cortex are shown. The inlay (grey background) shows images from an example mouse including photos of the head-plate surgery, as well as imaging frames from the WF-$Ca^{2+}$ imaging and EPI timeseries.

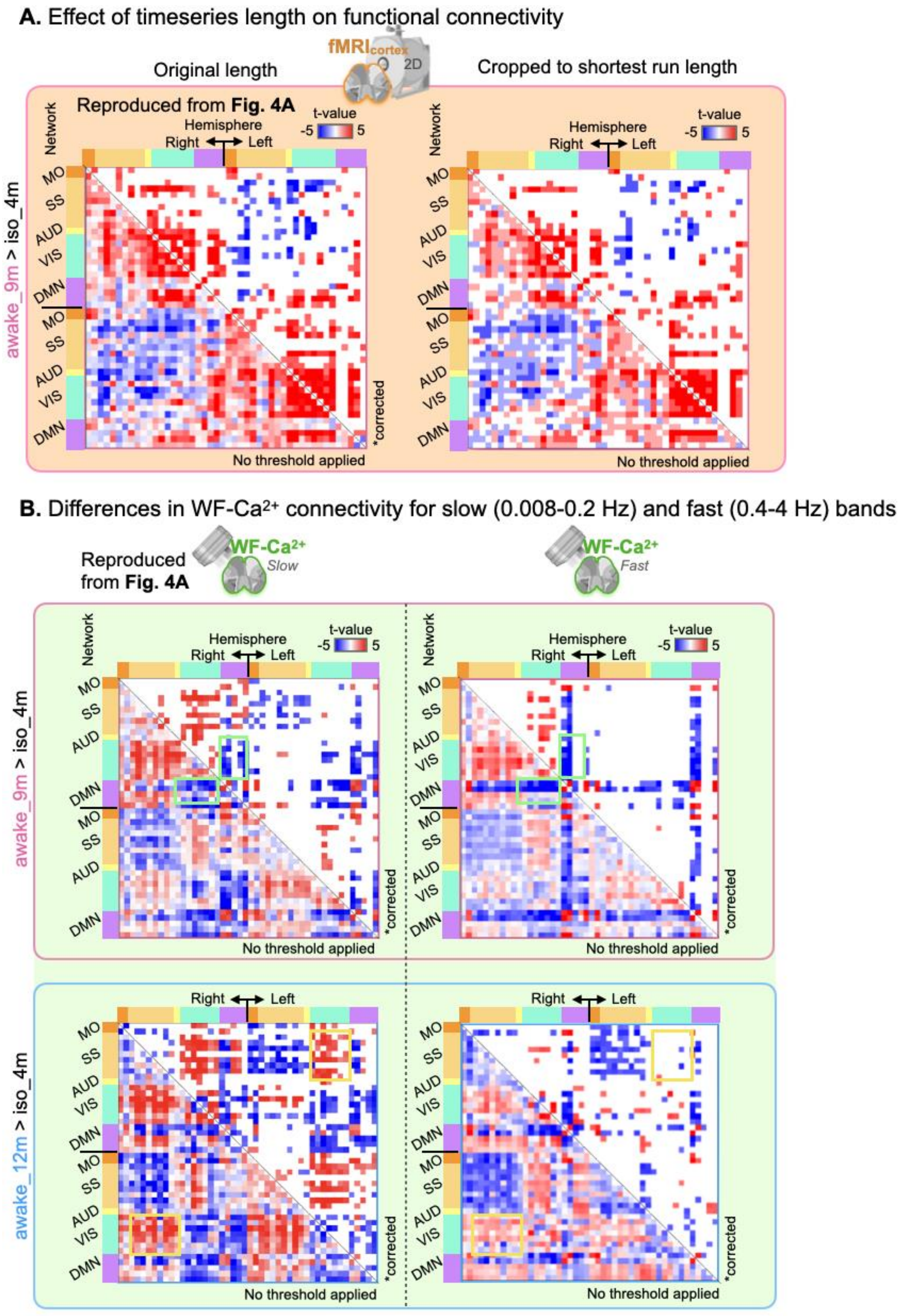


**Fig. S7. Differences in connectivity between anesthetized and awake mice: effects of timeseries lengths and frequency bands. A.** Left: Reproduction of **Fig. 4A**, fMRI results. Right: Differences in connectivity between awake and anesthetized sessions with timeseries truncated to the shortest retained scan length across all datasets (example for awake_9m vs. iso_4m). **B.** Left: Reproduction of **Fig. 4A**, WF-$Ca^{2+}$ results; Right: fast frequency band (0.4-4Hz) results.

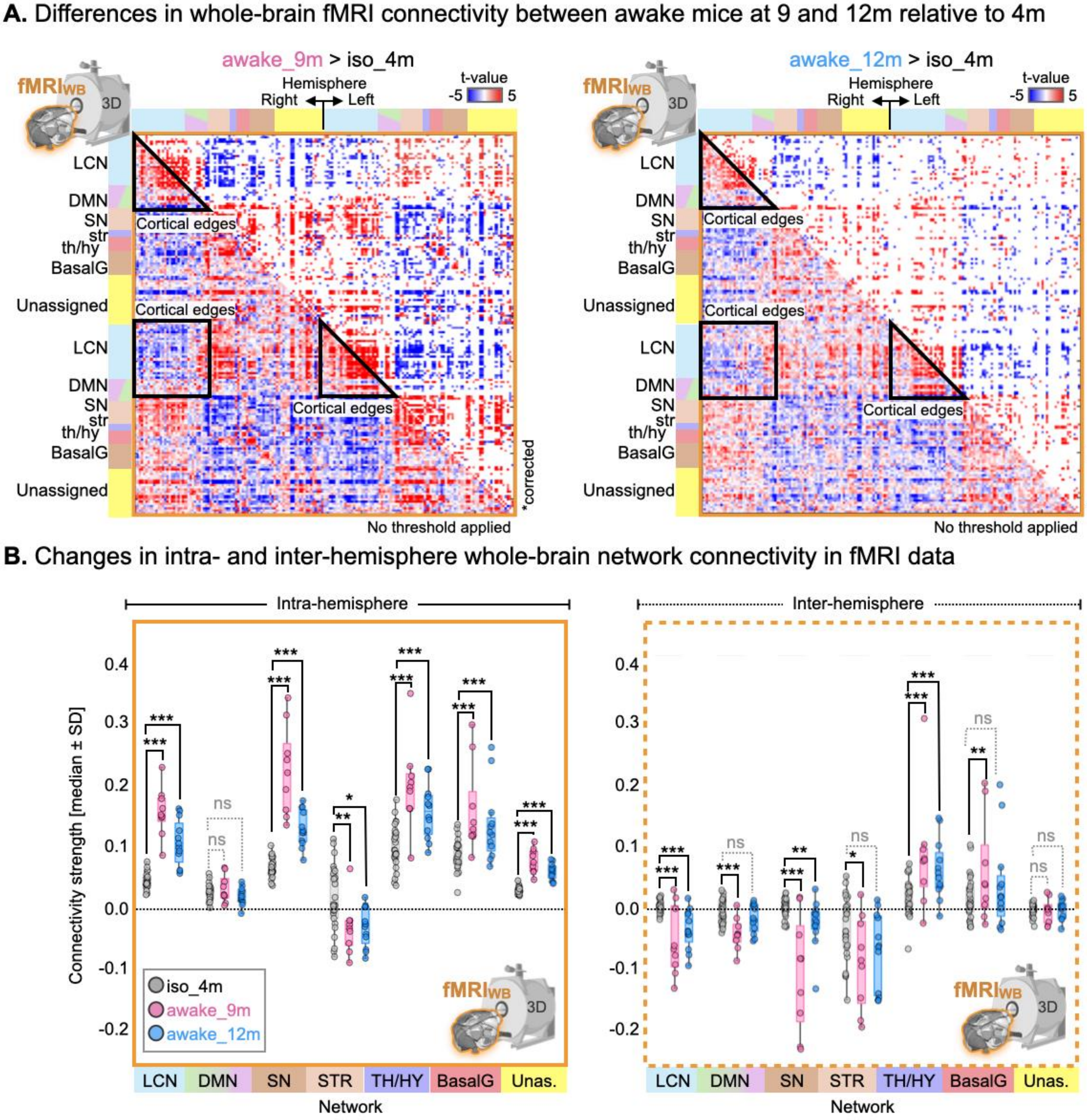


**Fig. S8. Differences in whole-brain fMRI connectivity between anesthetized and awake mice.** As-in **Fig. 4A & C** with fMRI data from the whole-brain and seven *a priori* networks, **Fig. S3**. Edges within the cortex are indicated in (**A.**) **(**black outlines).

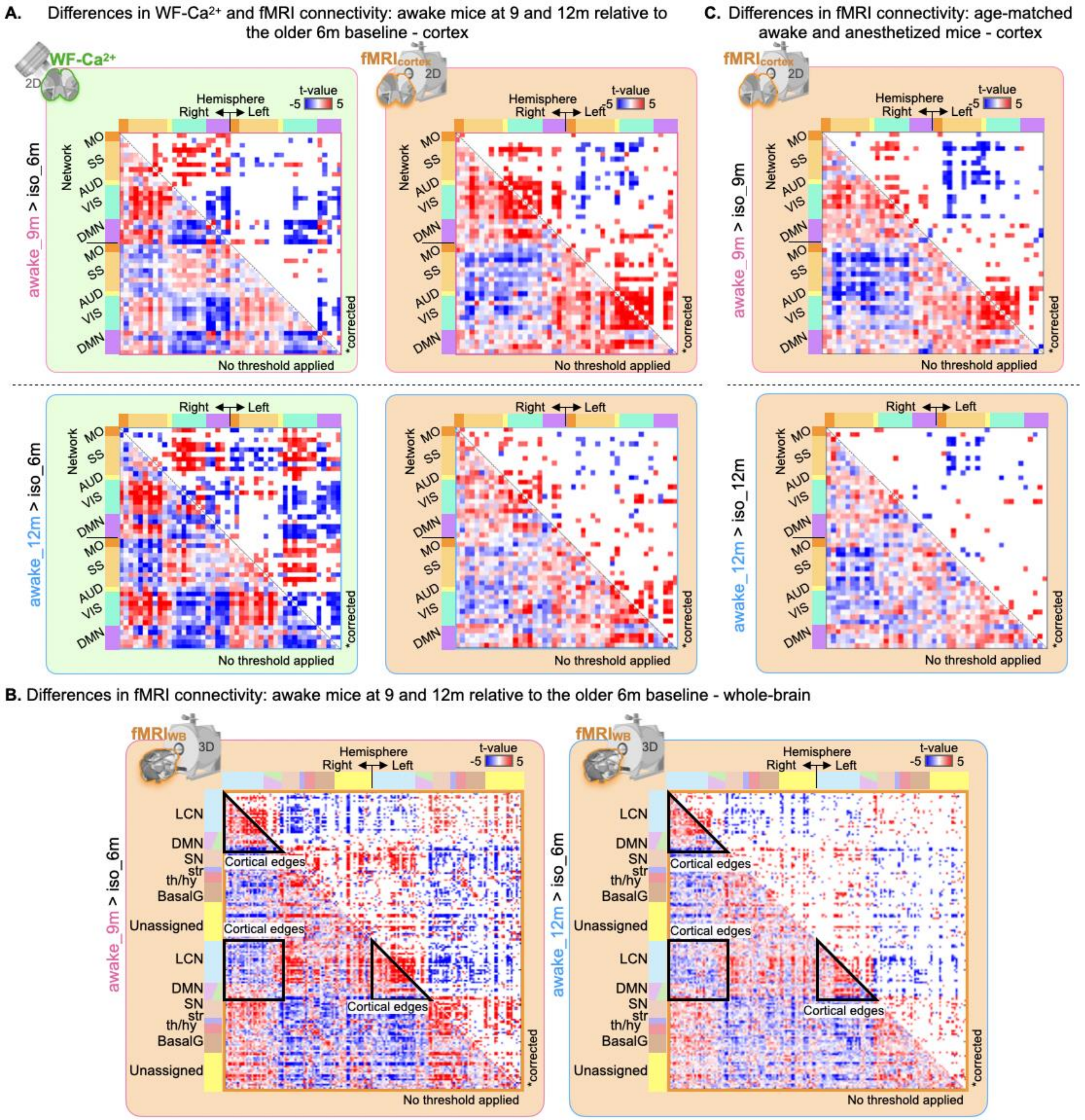


**Fig. S9. Differences in functional connectivity between awake mice at 9 and 12m relative to other (anesthetized) baseline imaging timepoints.** As-in **Fig. 3A** with data from the 6m (anesthetized) imaging timepoint used in-place of the 4m (anesthetized) imaging timepoint for cortical regions. Both WF-$Ca^{2+}$ imaging data (green panels, left) and fMRI data (orange panels, right) are included. **B.** As-in (**A.**) for whole-brain fMRI data. **C.** As-in (**A.**) using the age-matched anesthetized imaging timepoint (fMRI data only).

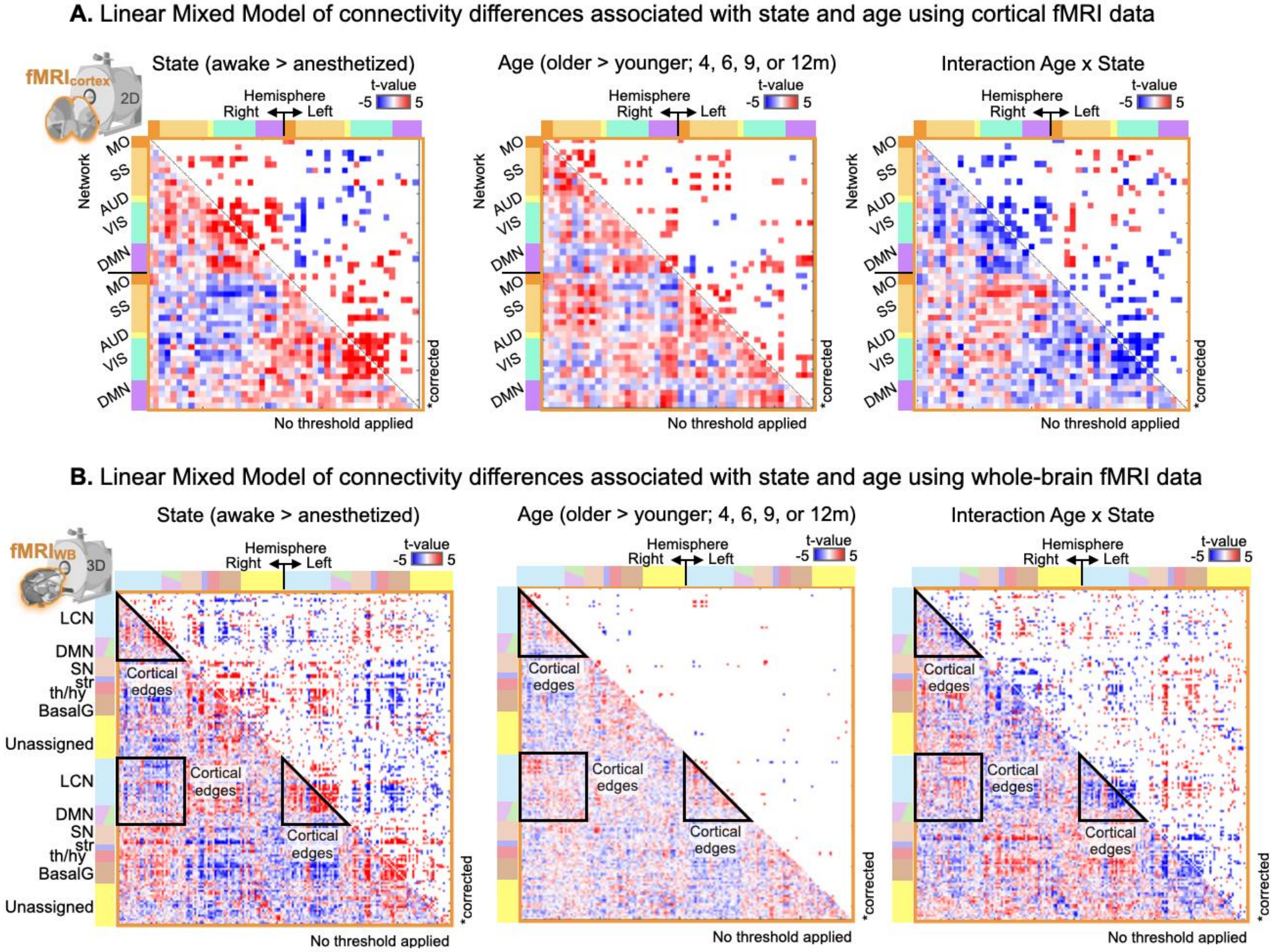


**Fig. S10. Results from two Linear Mixed Models built using fMRI data to assess contributions of age.** For each edge a linear mixed model tested the effects of State (awake versus anesthetized) and Age (4, 6, 9, or 12m) while accounting for animal identity as a random factor. The edges with significant effects (FDR-corrected, q<0.05) are displayed as t-value matrices for State (left), Age (middle) and their interaction (right). **A.** Model built using fMRI data from the cortex. **B.** Model built using fMRI data from the whole-brain. Edges within the cortex are indicated (black outlines).

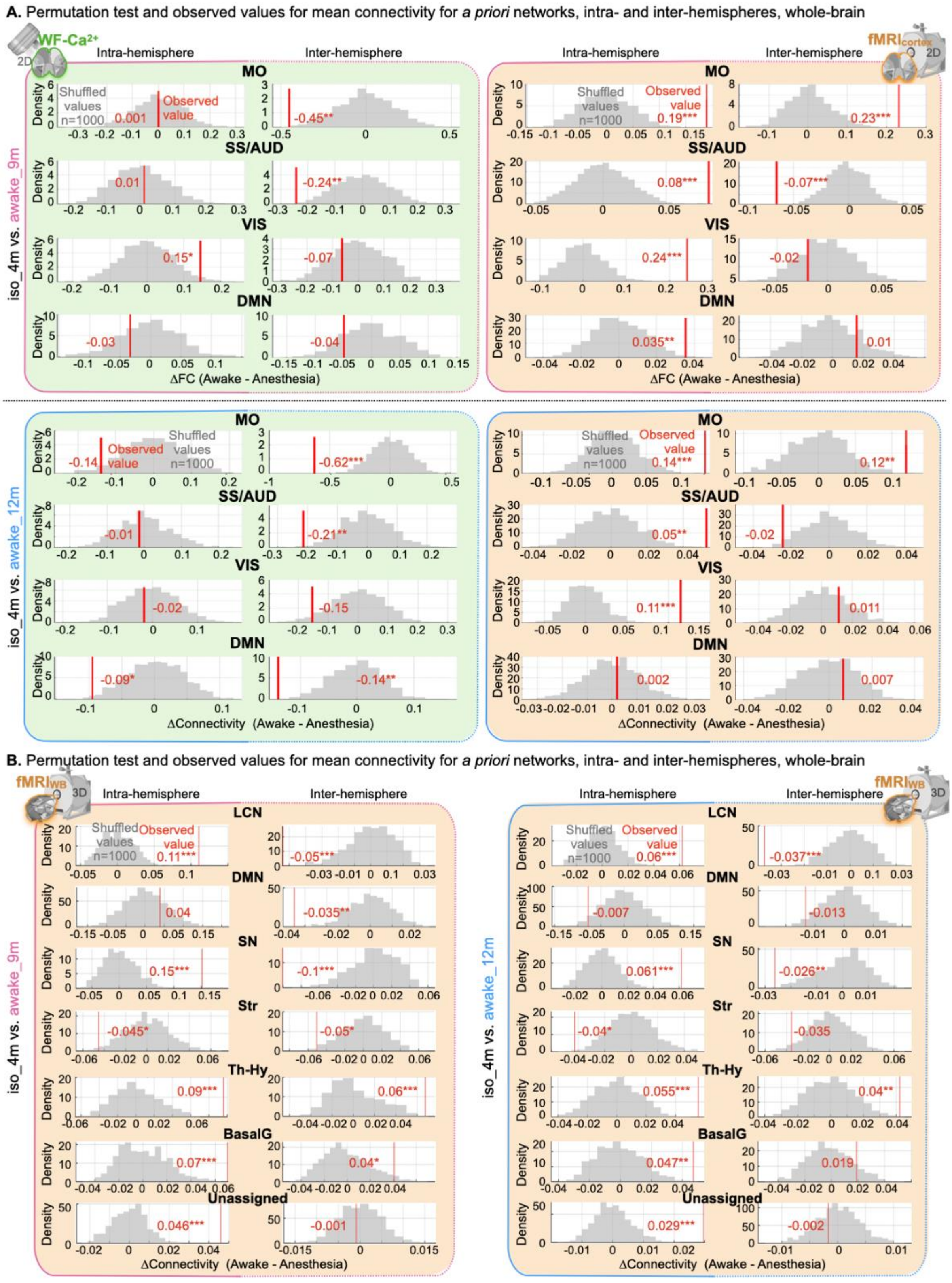


**Fig. S11. Permutation test results for multimodal data and fMRI data from the whole-brain.** For each network (rows) and hemisphere (columns: intra-hemisphere solid lines, inter-hemisphere dashed lines), histograms show mean connectivity strength differences generated from 1,000 permutations where awake and anesthetized labels were shuffled. ΔConnectivity plotted as t-values. **A.** Results from multimodal cortical data. **B.** Results from whole-brain fMRI data. Vertical red lines mark observed values from unshuffled data. Significance, corrected, displayed as: *p<0.05, **p<0.01, ***p<0.001).
